# Deep thermal proteome profiling for detection of proteoforms and drug sensitivity biomarkers

**DOI:** 10.1101/2022.06.10.495491

**Authors:** Nils Kurzawa, Matthias Stahl, Isabelle Leo, Elena Kunold, Isabelle Becher, Anastasia Audrey, Georgios Mermelekas, Wolfgang Huber, André Mateus, Mikhail M. Savitski, Rozbeh Jafari

**Author notes:** These senior authors contributed equally and both serve as corresponding authors. Correspondence should be addressed to M.M.S. and R.J. These first authors contributed equally to this paper (Nils Kurzawa, Matthias Stahl).

## Abstract

The complexity of the functional proteome extends significantly beyond the protein coding genome resulting in millions of proteoforms. Investigation of proteoforms and their functional roles is important to understand cellular physiology and its deregulation in diseases, but challenging to perform systematically. Here, we apply thermal proteome profiling with deep peptide coverage to detect functional proteoforms in acute lymphoblastic leukemia cell lines with different cytogenetic aberrations. We detect 15,846 proteoforms, capturing differently spliced, post-translationally modified, and cleaved proteins expressed from 9,290 genes. We identify differential coaggregation of proteoform pairs and establish links to disease biology. Moreover, we systematically make use of measured biophysical proteoform states to find specific biomarkers of drug sensitivity. Our approach thus provides a powerful and unique tool for systematic detection and functional annotation of proteoforms.

## Introduction

Proteins are the functional units expressed from genes and ultimately define the phenotype of cells. Through genomic variation (i.e. mutations and SNPs), alternative splicing of transcripts, proteolytic cleavage, post-translational modifications (e.g., phosphorylation, ubiquitination, acetylation and others), and protein-protein interactions (PPIs), the complexity of the functional proteome is expanded to millions of proteoforms ^1,2^. Therefore, identification and functional characterization of proteoforms can improve our understanding of biological processes in health and disease.

Although global proteoform measurement is critical for achieving full proteome characterization and annotation, its realization is still hampered by technological and analytical limitations. Top-down proteomics enables the precise characterization of proteoforms of individual proteins ^3^, and inference based on peptide level data from bottom up proteomics has recently been established ^4–6^. However, these approaches, while powerful, are either limited by proteome coverage, or by availability and variability of sample conditions which distinguish different proteoforms. Further, proteoforms have been detected representing protein sequence and post-translational modification status differences, but other important variations including protein complex and metabolite associations are difficult to distinguish without specific targeted experimental methods, and have therefore been excluded from identification. Recent initiatives have been proposed to define a human proteoform reference ^2,7^ and a reference map of proteoforms of human hematopoietic cells has recently been reported ^8^, and additional efforts are underway to address these gaps and improve knowledge of proteoforms.

Thermal proteome profiling (TPP) ^9,10^ is a method originally developed for unbiased detection of drug targets in living cells ^9,10^ and more recently tissues ^11,12^ by monitoring the changes in the thermal stability of proteins upon drug binding. It is implemented by applying the cellular thermal shift assay (CETSA) ^13,14^ on a proteome-wide scale using multiplexed quantitative mass spectrometry ^15^. Recent work has shown that TPP can not only inform on drug-target engagement ^16,17^ but also on protein-nucleic acid ^18^, protein-protein ^19^ and protein-metabolite interactions ^20^ as well as metabolic pathway activity ^21^ and the functional relevance of post-translational modifications ^22,23^. Moreover, it has been found that cell type-specific physiology is reflected in characteristic proteome thermal stability profiles and can be predictive of drug responses ^24^.

Here, we introduce the application of TPP for the detection of functional proteoforms. We demonstrate this by applying TPP without any perturbation to 20 different B-cell childhood acute lymphoblastic leukemia (cALL) cell lines, representing various disease subtypes defined by characteristic chromosomal rearrangements. In combination with high-resolution isoelectric focusing fractionation (HiRIEF) ^25^ we measure thermal stability with unprecedented peptide coverage per gene. This aspect is exploited to infer functionally relevant proteoforms in an unbiased manner, capturing differently spliced, modified or cleaved proteins expressed from the same gene. We link differentially thermally stable proteoforms across cell lines with the developmental stage of the cell of origin and the genetic subtypes of the cALL samples. Moreover, we analyze differential coaggregation of pairs of proteoforms across the different cALL cell lines and link coaggregation to disease biology. Lastly, we systematically make use of measured biophysical proteoform states to find biomarkers for cell line sensitivity to 528 oncology and investigational compounds.

## Results

### Deep thermal profiling across cALL cell lines allows unbiased assignment of peptides to functional proteoforms

To systematically measure the melting behavior of proteins in cALL cell lines representing different molecular subtypes, we performed temperature-range TPP ^9^ with 8 temperatures per sample and multiplexed two cell lines at a time using TMTpro ^26,27^ (Supplementary Data 1). We profiled cell lines that reflect different cALL subtypes, as defined by diverse genomic rearrangements, a balanced mix of female and male donor patients and different B-cell developmental stages of origin (Fig. 1a). We obtained deep peptide coverage per gene symbol (Supplementary Figure 2a) by measuring a total of 114 HiRIEF fractions per sample before LC-MS/MS analysis ^25^. In total, we identified 243,929 unique peptides mapping to 16,094 gene symbols across cell lines with comparable global melting profiles (Fig. 1b).

**Figure 1:**
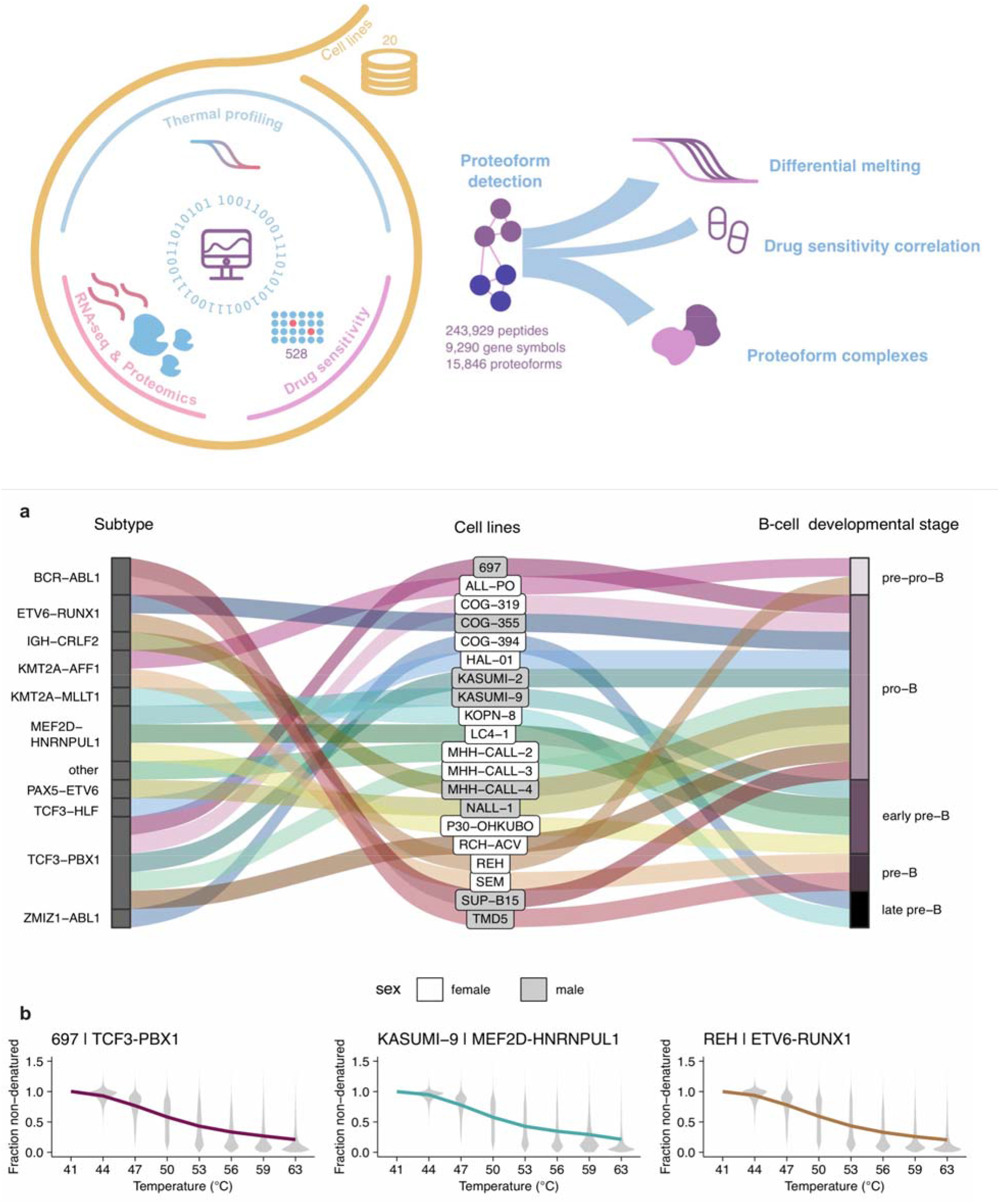
a) Alluvial diagram representing profiled cell lines and their characteristics. b) Exemplary average melting profiles across all peptides identified and quantified in the cell lines 697, KASUMI-9 and REH after normalization.

**Figure 2:**
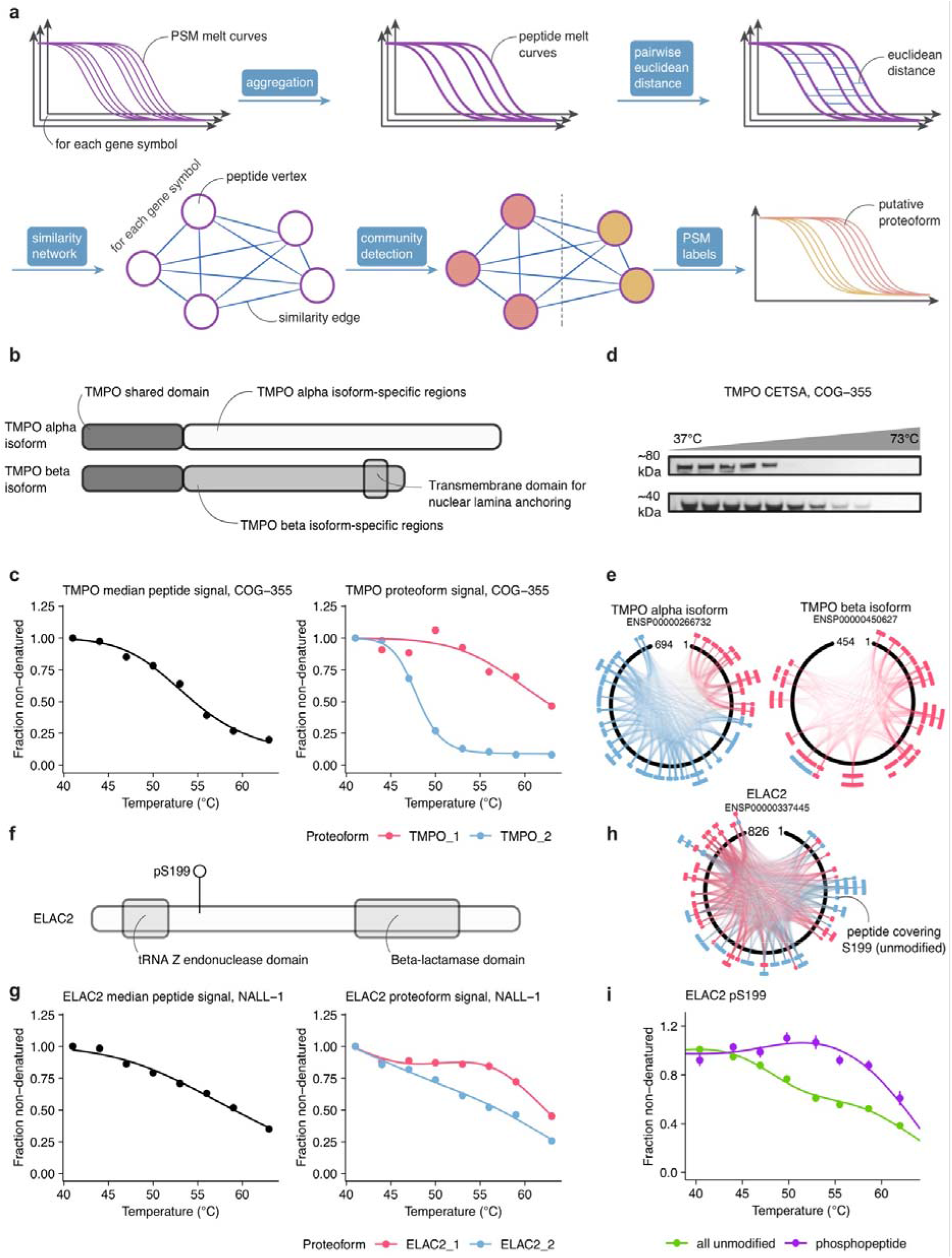
Proteoform detection based on peptide melting curve similarity. a) Schematic of the method. b) Schematic of the domains of the TMPO alpha and beta isoforms. c) Melting curves of the median TMPO peptide signal (left) and by proteoforms as detected by the above outlined approach. d) Western blot validation of the detected proteoforms for TMPO (alpha on top, beta below) with differential thermal stability in the COG-355 cell line. e) Mapping of peptides assigned to the different proteoforms to the alpha and beta isoforms of TMPO. f) Schematic of the protein domains of ELAC2. g) Melting curves of the median ELAC2 peptide signal (left) and by detected proteoforms. h) Mapping of peptides assigned to the different ELAC2 proteoforms to the protein sequence. i) Melting curves of unmodified and pS199 phosphorylated ELAC2 in HeLa cells from Potel et al. (2021).

As expected for proteoforms with different cellular functions, we observed that peptides mapping to a single gene symbol often formed groups with distinct thermal stability patterns. In fact, grouping of peptides by thermal stability reflected annotated proteoforms to some extent (Supplementary Figure 2b,c). We thus exploited clustering of similar peptide melting profiles by developing a method to assign peptides to different proteoforms without relying on their annotation (Fig. 2a). To do so, we filtered our dataset to contain only peptides that had been identified and quantified in at least two cell lines and computed pairwise similarities between all melting curves of peptides mapping to the same gene symbol. Then, for each gene symbol, a fully connected graph was constructed based on respective peptide similarities, and clusters were detected using the Leiden algorithm ^28^. We accepted all recovered clusters supported by at least five unique peptides and modularity *Q >* 0, indicating that the magnitude of division of the graph network into two or more clusters was higher than expected by chance, and assigned them to different proteoform IDs. This resulted in detection of 15,846 proteoforms of 9,290 genes, with the majority of genes being represented by one (44%) or two (44%) and a maximum of five proteoforms (Supplementary Figure 3a, Supplementary Data 2). As expected, our derived proteoforms showed higher modularity than ENSEMBL annotated ones (Supplementary Figure 3b), suggesting that this approach extends delineation of proteoforms in comparison to existing annotations.

**Figure 3:**
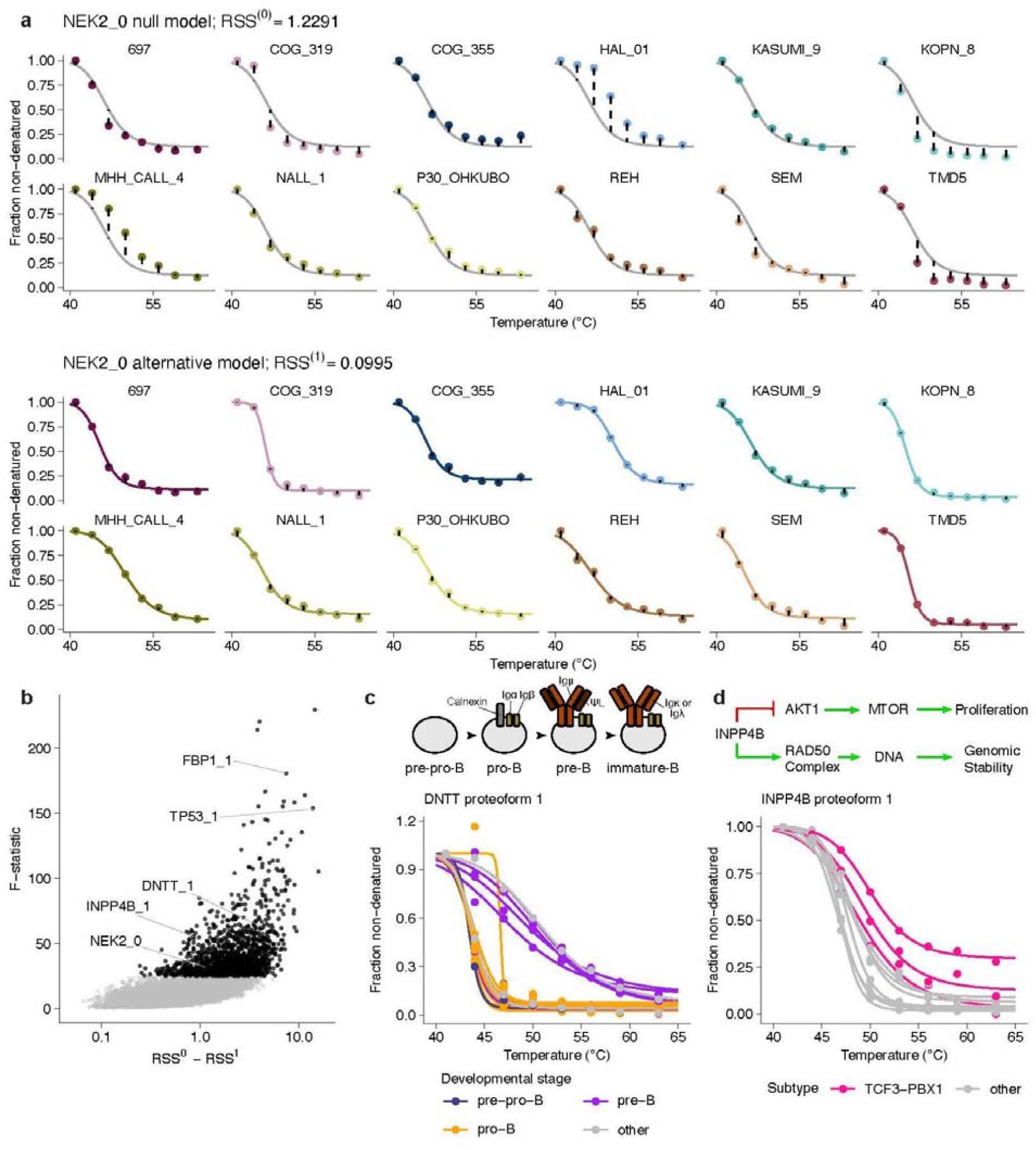
Nonparametric differential analysis of melting curves (NPARC) across cell lines. a) Exemplary fits of the null (top) and alternative (bottom) models to the cell line melting profiles of NEK2_0, the only proteoform found for NEK2. b) Volcano plot of the results obtained from the differential analysis. Black points represent proteoforms with the top 10% of *F*-statistics taken as differentially melting across cell lines (*F* _≥_*P*_90%_(F)). c) Melting profile of DNTT proteoform 1 (DNTT_1) across cell lines color labeled by B-cell developmental stage. d) Melting profile of INPP4B proteoform 1 (INPP4B_1) across cell lines color labeled by cell line genomic aberration subtype.

When examining detected proteoforms in detail, we confirmed our approach by identifying proteoforms representing previously described cases of alternative splicing and proteolytic cleavage. For example, lamina-associated polypeptide 2 (TMPO) is a protein known to be expressed in several isoforms generated via alternative splicing. Two functionally important isoforms, alpha and beta, share a common N-terminus, but differ in their C-termini ^29^. The TMPO alpha isoform associates with chromatin in a cell cycle-dependent manner, and TMPO beta isoform which associates with the inner nuclear lamina via a transmembrane domain and facilitates lamin-mediated structural organization of chromatin (Fig. 2b) ^30^. Using our proteoform detection method, we found two distinctly melting proteoforms for TMPO (Fig. 2c). We used an antibody recognizing the TMPO N-terminus to confirm differential melting for bands at molecular weights corresponding to alpha and beta isoforms (Fig. 2d). Furthermore, when inspecting the underlying peptides in terms of their mapping to the TMPO ENSEMBL isoforms, we observed that the majority of peptides assigned to proteoform 1 (TMPO_1) were either specifically mapping to the sequence of the TMPO beta isoform or to the joint N-terminus of both isoforms (Fig. 2e). Thus, our method successfully detected the TMPO alpha and beta isoforms solely by considering the melting profiles of the peptides across cell lines mapping to the respective gene symbol.

In another example, we identified two proteoforms of the Zinc phosphodiesterase ELAC2 (Fig. 2f), an enzyme known to localize to the nucleus and to mitochondria ^31^. While ELAC2_2, comprising an unmodified peptide covering serine 199 (S199), showed a profile similar to the median peptide signal per gene symbol, ELAC2_1 displayed a pattern (Fig. 2g) reminiscent of a differentially melting proteoform phosphorylated on S199 that we observed in a previous study using a phosphoTPP experiment ^22^. To corroborate ELAC2_1 as pS199 phospho-proteoform of ELAC2, we queried our dataset against the human database, this time including phosphorylation as a modification. In fact, we found a peptide capturing the pS199 site of ELAC2 that, when inspected for thermal stability, showed a pattern similar to ELAC2_1 and the pS199 phospho-proteoform identified in the phosphoTPP experiment (Supplementary Figure 4). Therefore, our proteoform detection approach successfully identified post-translationally modified subpools of the same protein without the need for peptide enrichment.

**Figure 4:**
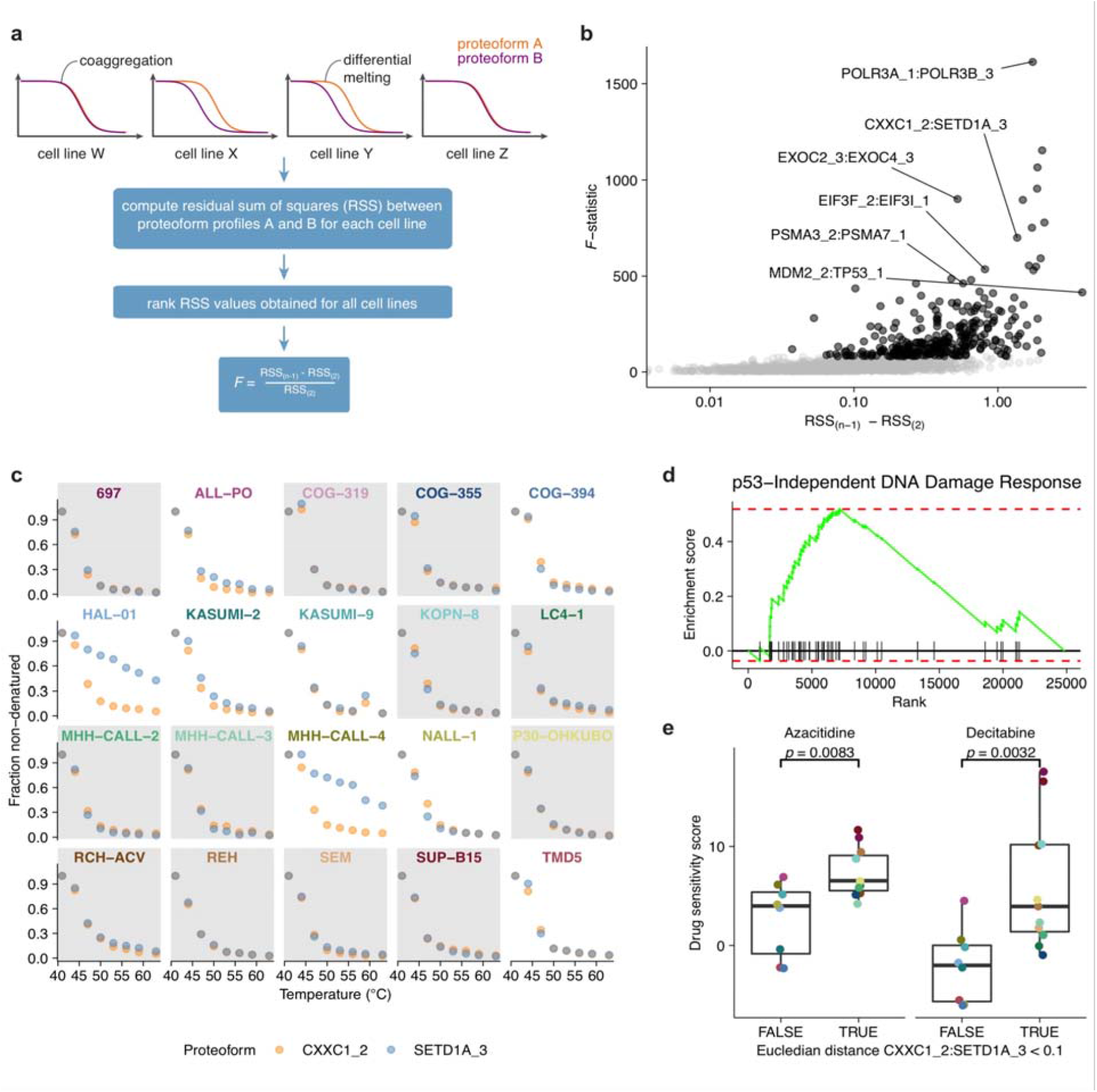
Differential proteoform coaggregation analysis. a) Schematic of the performed analysis. To obtain robust results, the *F*-statistic is computed based on the difference of the second highest (RSS_(n-1)_) and second lowest (RSS_(2)_) residual sum of squares between the two proteoforms in all cell lines. b) Volcano plot of the results of the analysis. RSS_(n-1)_ - RSS_(2)_ represents the effect size, i.e. the difference between the profiles of proteoform A and B in the cell line with the second highest and second lowest distance. c) Profiles of CXXC1_2 and SETD1A_3 across cell lines showing their coaggregation in some (gray background) and differential melting (white background) in other cell lines. d) Enrichment plot for genes part of the “p53-Independent DNA Damage Response” set based on differentially expressed transcripts between cell lines with coaggregation of CXXC1_2 and SETD1A_3 vs. all others (NES = 1.82; *p*_adj._ = 0.01). e) Boxplots of drug responses of cell lines with CXXC1_2 and SETD1A_3 coaggregation vs. all others to two different nucleoside analogs. The p-values shown were obtained from a two-sided Welch two-sample t-test. Center lines in all box plots represent the median, the bounds of the boxes are the 75 and 25% percentiles i.e., the interquartile range (IQR) and the whiskers correspond to the highest or lowest respective value.

In addition to these examples of alternatively spliced isoforms and post-translationally modified proteoforms, we found several cases of proteoforms that resulted from proteolytic cleavage, e.g. Pre-saposin (PSAP) (Supplementary Figure 5a-d) and NOTCH1 (Supplementary Figure 5e-g) These results are also in agreement with previous studies that established the existence and biological relevance of these proteoforms, further validating our approach.

**Figure 5:**
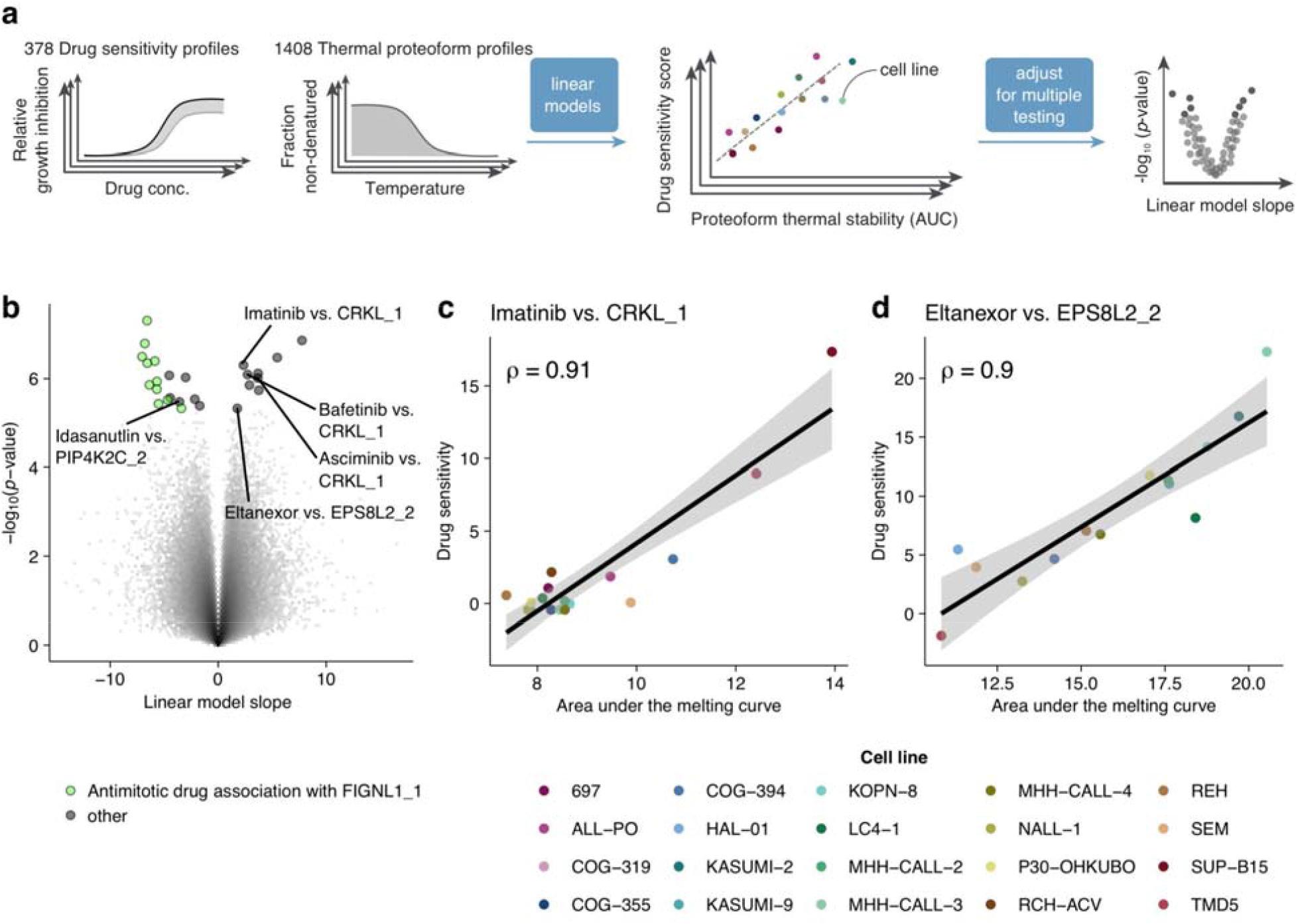
Association of thermal proteoform stability to drug viability across cALL cell lines. a) Schematic of the strategy to test for thermal proteoform stability drug associations. b) Volcano plot representing the results obtained from the analysis depicted in a). c,d) Scatterplot of CRKL_1 and EPS8L2_2 thermal stability and cell line sensitivity to Imatinib and Eltanexor, respectively.

Taken together, the peptide-level TPP data and the new proteoform detection algorithm allowed us to identify different proteoforms that reflected known functional characteristics of the respective proteins.

### Differential thermal proteoform stability across samples reflects B-cell precursor origins and disease subtypes

Thermal stability of proteins can vary across cell lines, reflecting genomic variation, specific protein-interactions networks and differential pathway activity ^24^. To explore this aspect we sought to identify differential thermally stable proteoforms across our samples. In order to do so, we performed Nonparametric Analysis of Response Curves (NPARC) ^32^ to find differences across the 20 cALL cell lines (illustrated by NEK kinase peptide profiles in Fig. 3a). This allowed us to detect 1408 proteoforms with differential melting curves (90% percentile of observed *F*-statistics) across the profiled cell lines (Fig. 3b, Supplementary Data 3). Among the top hits of the analysis we found a proteoform of p53 (TP53_1), a tumor suppressor protein and fructose-1,6-bisphosphatase 1 (FBP1_1), a rate-limiting enzyme of gluconeogenesis (Supplementary Figure 6a). For the NPARC hits, we sought to identify potential mechanisms behind differential melting of these proteoforms, and therefore annotate differences in proteoform functional roles in different cell line backgrounds. While the differential thermal stability of TP53_1 could be related to altered protein interactions (see next section), in the case of FBP1_1 the higher thermal stability of the cluster of proteoform peptides was associated with high FBP1 protein abundance in respective cell lines (*p* = 3.3e-10, two-sided Welch two sample t-test on protein fold changes, Supplementary figure 6b). However, as we did not observe global correlation between thermal stability and abundance (Supplementary figure 6c), these data suggested a specific effect linked to higher FBP1 activity in these cell lines. Previous studies had shown that cell lines with high FBP1 abundance display activation of the pentose phosphate pathway resulting in chemotherapy resistance and poor clinical outcome in acute myeloid leukemia ^33^. In agreement with these observations, we found higher thermal stability of all proteoforms of glucose-6-phosphate dehydrogenase (G6PD), the rate-limiting enzyme in the oxidative pentose phosphate pathway, in the cell lines with high thermal stability of FBP1_1 (Supplementary Figure 6d). This illustrates how our data can be used to identify functional links based on the differential thermal stability of proteoforms.

Another protein with differential thermal stability across cell lines was the DNA nucleotidylexotransferase (DNTT_1), a DNA polymerase which adds random nucleotides to the junction of rearranged immunoglobulin chains during B-cell maturation ^34^. We identified strikingly distinct melting profiles (Fig. 3c) that were associated (*p* = 0.036, Fisher test) with the developmental stage of the B-cell progenitor cell of origin of the acute lymphoblastic leukemia lines ^35^. DNTT diversifies the variable region of the Ig-light chain during the pre-B stage, and diversifies Ig-heavy chain variable regions during the pro-B stage ^36,37^. Thus, higher thermal stability may indicate differences in DNTT-DNA binding dynamics between the developmental stages.

We also found that INPP4B_1, a proteoform of INPP4B, a protein and lipid phosphatase that antagonizes the PI3K/Akt signaling pathway ^38^, showed higher thermal stability in cell lines of the TCF3-PBX1 subtype (*p* = 0.017, two-sided Wilcoxon rank sum test on area under the melting curves). INPP4B has also been shown to be involved in maintaining genomic integrity through associations with RAD50 in the nucleus, and loss of INPP4B was shown to sensitize cells to PARP inhibition ^39^. We observed that TCF3-PBX1 fusion cells had decreased INPP4B abundance at baseline ^35^ (*p* = 0.039, two-sided Welch two sample t-test on protein fold changes) and were selectively sensitive to the PARP inhibitor Talazoparib (*p* = 0.011, two sided Welch two sample t-test on sDSS scores). This suggests that this proteoform is associated with reduced genomic stability, which could implicate nuclear relocalization in the cell lines with observed high thermal stability.

Overall, we detected hundreds of examples of proteoforms with differential thermal stability in the cALL cell lines studied. Since thermal stability reflects the state and activity of proteins in a complementary way to traditional abundance proteomics ^18,21^, these examples pinpoint pathway activation status and reveal new candidate biomarkers for therapy.

### Proteoform coaggregation analysis reveals differential protein-protein interactions across cell lines

Melting curves of interacting proteins (protein-protein interactions, PPI) or complex members have been shown to often coincide, a feature attributed to coaggregation of the respective interactors ^19^. Recently, we have exploited this concept to test for differential coaggregation of protein interactors between two conditions ^40^. Here, we adapted this approach to a robust multi-group comparison (Fig. 4a) to detect differential proteoform-proteoform interactions (PFPFI). across the profiled cALL cell lines using an extended PPI annotation of STRING-database ^41^. In total we tested 2,901 PFPFI, which showed coaggregation in at least one of the cell lines, for differential coaggregation across cell lines. We considered PFPFI within the top 10% of obtained *F*-statistics (290 PFPFIs) as significantly differential across cell lines (Fig 4b, Supplementary Data 4). Among those cases we identified several examples of differential intra-complex PFPFIs, potentially reflecting varying degrees of complex assembly or activity across the profiled cell lines (Supplementary Figure 7a-d).

One differentially coaggregating proteoform pair was MDM2_2 and TP53_1 (Supplementary Figure 7e). MDM2 is an E3 ubiquitin ligase which is known to ubiquitinate the tumor suppressor p53 and thus promote its degradation ^42^. Furthermore, MDM2 is often upregulated in different cancers leading to increased degradation of p53, resulting in uncontrolled cell division ^43^. We thus wondered whether cell lines in which MDM2_2 and TP53_1 coaggregated, which we interpreted as a sign of MDM2 binding to p53 and thus promoting its degradation, were more susceptible to MDM2 inhibition than other cell lines. Indeed, the two cell lines which featured coaggregation of MDM2_2 and TP53_1, LC4-1 and P30-OHKUBO, showed higher sensitivity to Idasanutlin, an MDM2 inhibitor, compared to other cell lines (*p* = 0.02, two-sided Welch two sample t-test, Supplementary Figure 7f). This showcases how our strategy can reveal functionally relevant connections between proteins and use them to generate hypotheses on drug sensitivity.

We also found the differentially coaggregating proteoform pair CXXC1_2 and SETD1A_3 (Fig 4c). SETD1A is a SET domain containing histone methyltransferase which has been reported to mediate DNA damage response ^44^ and CXXC1 was found to regulate SETD1A activity ^45^. We hypothesized that coaggregation of CXXC1_2 and SETD1A_3 could reflect an ongoing DNA damage response in respective cell lines. In fact, comparing RNA-seq profiles ^35^ of cell lines with coaggregating versus differential CXXC1_2 and SETD1A_3 melting profiles revealed that the gene set “p53-Independent DNA Damage Response” was significantly enriched among upregulated genes in cell lines which featured coaggregation of this proteoform pair (Fig. 4d). We further asked whether these cell lines showed altered sensitivity to DNA damage inducing drugs, such as nucleoside analogs. Consistent with this hypothesis, we observed significantly higher sensitivity to the nucleoside analogs and hypomethylating agents azacitidine and decitabine for cell lines in which CXXC1_2 and SETD1A_3 coaggregated (Fig. 4e).

Taken together, we present an approach for the detection of differentially coaggregating pairs of proteoforms and show that some of these altered interactions can be linked to activity of cellular processes and drug response.

### Systematic evaluation of proteoform thermal stabilities as biomarkers for drug response

Encouraged by the observed associations between pathway activity (reflected in protein thermal stability) and drug sensitivity, we sought to generalize this principle across a larger drug panel, namely the 528 drugs used in our previous study ^35^. By using *limma* ^46^ to correlate drug sensitivity scores (DSS) of 378 drugs with a minimal effect cutoff on any of the profiled cell lines (DSS >= 6) with all previously determined 1408 differentially thermally stable proteoforms (Fig. 5a), we retrieved 26 significant drug-proteoform thermal stability associations (*p*_*adj*._ *<* 0.1, Benjamini-Hochberg method) (Fig. 5b, Supplementary Figure 5). Among these, we found thermal stability of CRKL_1 to be positively correlated with sensitivity to the BCR-ABL inhibitors Imatinib, Asciminib and Bafetinib (Fig. 5c). CRKL is an adapter protein downstream of ABL1 that is phosphorylated upon activation of ABL1 ^47^. Previously it was observed that CRKL was thermally destabilized upon treatment with dasatinib, another BCR-ABL inhibitor ^9^. Inversely, thermal stabilization of CRKL appears to be related to active ABL1 signaling which is in line with a positive correlation of sensitivity to BCR-ABL1 inhibitors (Supplementary Figure 8a). Moreover, we found that cell line sensitivity to several antimitotic drugs was negatively correlated with Figetin-like protein (FIGNL1) proteoform (FIGNL1_1) thermal stability (Fig. 6e, Supplementary Figure 9a-f). FIGNL1 is involved in DNA double strand repair via homologous recombination^48^. Since FIGNL1_1 thermal stability was negatively correlated with FIGNL1 protein abundance (Supplementary Figure 9g), high FIGNL1_1 thermal stability could reflect active engagement in the FIGNL1-containing complex to resolve DNA double strand breaks. Indeed, correlation of FIGNL1_1 thermal stability with antimitotic drug sensitivity was stronger than for FIGNL1 abundance (Supplementary Figures 9h and i). Thus, high activity of the FIGNL1-containing complex could lead to reduced mitotic exit at cell cycle checkpoints and may thus explain lower sensitivity to antimitotic drugs.

Another interesting hit was the positive correlation of PIP4K2C_2 thermal stability with cell line sensitivity to the MDM2 inhibitor Idasanutlin (Fig. 6e, Supplementary Figure 10a and b). Several PIP4K2 family members have previously been linked to promotion of tumorigenesis in the context of p53 loss of function ^49^. Thus, it appears plausible that high thermal stability of PIP4K2C_2, potentially reflecting a higher fraction of cofactor-bound protein pool, is associated with increased sensitivity to MDM2 inhibition, as the related signaling pathway appears to only lead to cell growth in the absence of p53 function. The fact that the correlation of Idasanutlin sensitivity to PIP4K2C abundance is weaker and positive rather than negative (Supplementary Figure 10c), reinforces the notion that thermal stability gives a more functional readout of protein state compared to measurements of protein abundance.

Finally, we detected a positive correlation between EPS8L2_2 (Epidermal growth factor receptor kinase substrate 8-like protein 2) with Eltanexor (Fig. 5c), a nuclear export inhibitor. EPS8L2 is known to form a complex with SOS1 and ABI1 which is involved in regulating actin remodeling ^50^. The observed correlation was specific to thermal stability and to the EPS8L2_2 proteoform (Supplementary Figure 11a-c). To investigate how high EPS8L2_2 thermal stability could confer sensitivity to nuclear export inhibition, we performed a differential expression analysis between cell lines with high and low EPS8L2_2 thermal stability. When performing GO-molecular function enrichment analysis on the transcripts up-regulated in cell lines with high EPS8L2_2 thermal stability we found a significant enrichment (*p*_adj._ < 0.1) of the terms ‘actin binding’, ‘antigen binding’ and ‘immunoglobulin receptor binding’. This may indicate that high EPS8L2_2 could reflect actin remodeling in response to B-cell receptor (BCR) activation ^51^. It has been shown previously that nuclear export inhibition suppresses downstream effects of BCR signaling in chronic lymphocytic leukemia ^52^, therefore it is plausible that Eltanexor treatment may be effective in a subset of acute lymphocytic leukemias relying on BCR signaling for proliferation.

## Discussion

CETSA and TPP were developed with the primary goal of detecting protein targets of drugs ^9,14^. However, it has been realized that these methods can also detect other sources of protein biophysical variation which are difficult to quantify with other proteomics methods, including protein interactions with other biomolecules ^53^. Since the adaptation of the method to infer functional phosphorylation sites ^22,23^, it has also become clear that TPP bears the potential for detecting post-translationally modified proteoforms. Here we have performed TPP with unprecedented peptide coverage and generalized this concept to enable unbiased detection of co-existing functional proteoforms. The detected events of diversified protein products comprise cases of alternative splicing, proteolytic cleavage, post-translational modifications, and variants interacting with metabolites, proteins, or DNA. Previous efforts to detect proteoforms have not been able to distinguish between entities representing these sources of variation in a global and unbiased way.

We show that performing TPP with high peptide coverage allows for the detection of proteoforms and simultaneously allows inference of functional aspects by revealing peptide sequence coverage, differences in proteoform-proteoform interactions, and associations to drug response. By integrating thermal stability of proteoforms, transcriptomics and drug sensitivity profiling data across cell lines we demonstrate that it is possible to identify biomarkers for cellular processes and drug response. Thus, we believe that the broadly applicable deep thermal proteome profiling for proteoform detection is a powerful and complementary addition to existing technologies for delineating proteoforms and for supporting analytical strategies interrogating cellular processes at the molecular level.

## Methods

### Cell cultivation

The 20 childhood B-cell Precursor Acute Lymphoblastic Leukemia (BCP-ALL) cell lines used in this study were obtained from Deutsche Sammlung von Mikroorganismen und Zellkulturen GmbH (DSMZ, German Collection of Microorganisms and Cell Cultures, Braunschweig, Germany), from Children’s Oncology Group ^54^ Childhood Cancer Repository (Lubbock, TX, USA), American Type Culture Collection (ATCC), Japanese Collection of Research Bioresources Cell Bank (JCRB), European Collection of Authenticated Cell Cultures (ECACC, England), and Banca Biologica e Cell Factory (San Martino, Italy). Roswell Park Memorial Institute (RPMI) 1640 (AQmedia, Sigma-Aldrich) or Iscove’s Modified Dulbecco’s Medium (IMDM, Sigma-Aldrich) supplemented with either 10% or 20% fetal bovine serum (FBS, Sigma-Aldrich), 20 mM HEPES (Gibco/Life Technologies), 1 mM sodium pyruvate (Sigma-Aldrich), 1x MEM non-essential amino acids (Sigma-Aldrich), and 1x Penicillin-Streptomycin (Sigma-Aldrich) was preferably used. Cell line provider details, culture conditions, and growth media are also described in supplementary data 1 of ^35^. Cell lines were grown at 37°C and 5% CO2 to a cell density of approximately 1-2 million cells/mL. Cells were harvested at 500 x g for 3 min and washed twice with Hank’s Balanced Salt Solution (Gibco™ HBSS, no calcium, no magnesium, no phenol red).

#### Sample preparation for LC-MS/MS

### Thermal proteome profiling of the cell lines

Freshly washed cells were resuspended to a density of 100 million cells/mL in HBSS and distributed as aliquots of 10 million cells into eight 0.2-mL PCR tubes. Tubes were heated in parallel for 3 min to 41, 44, 47, 50, 53, 56, 59 and 63°C, followed by a 3-min incubation time at room temperature. Afterwards, cells were flash-frozen in liquid nitrogen.

### Digest and TMT labeling

Lysis was performed by five freeze-thawing cycles using a 25°C heating block and liquid nitrogen. Cell debris and precipitated proteins were removed by centrifugation at 21000 x *g* and 4°C for 40 min. Supernatants were transferred to new tubes and protein concentrations were determined using the DC protein assay according to standard protocols provided by the kit manufacturer (Bio-Rad, Hercules, CA). Equal volumes of soluble protein supernatants were transferred to new tubes and subjected to in solution digestion. First, the samples were supplemented with reagents to contain a final concentration of 50 mM TEAB, 0.1% SDS and 5 mM TCEP. Reduction was performed at 65°C for 30 min. Samples were then cooled down to RT and alkylated with 15 mM of chloroacetamide for 30 min. Proteins were digested overnight with 1 to 40 Lys-C (Wako Chemicals GmbH, Neuss, Germany)-to-protein ratio. Consecutively, trypsin (Thermo Fisher Scientific, Waltham, MA) was added at a 1 to 70 enzyme-to-protein ratio for an eight hours incubation at 37°C. Finally, the same amount of trypsin was added one more time for a second overnight incubation. Resulting peptides were labeled by 16-plex TMTpro-tags (TMTpro, Thermo Fisher Scientific, Waltham, MA, USA) using the same amount of respective label for each sample. Eight melting points of two randomly selected cell lines were combined in each TMT 16-plex set. The protein amounts were adjusted to contain the same total protein amount for all cell lines throughout the TMT sets. An overview of the sets is given in Supplementary Data 1. Labeling was performed according to manufacturer’s instructions but with 2 h instead of 1 h incubation prior to quenching the TMT labeling reaction. Labeling efficiency was determined by LC-MS/MS before mixing the TMT labeled samples. For sample clean-up solid phase extraction using SPE strata-X-C columns (Phenomenex, Torrance, CA, USA) was performed. Purified peptides were finally dried in a vacuum centrifuge.

### High resolution isoelectric focusing (HiRIEF) of peptides

The prefractionation method was applied as previously described ^25^. Sample pools of ∼300 μg were subjected to peptide IEF-IPG (isoelectric focusing by immobilized pH gradient) in pI range 3 - 10 and 3.7 - 4.9 respectively. Dried peptide samples were dissolved in 250 μL rehydration solution of 8 M urea containing 1% IPG pharmalyte pH 3 – 10 or 2.5 – 5, respectively (GE Healthcare) and allowed to adsorb to the gel bridge strip and the 24 cm linear gradient IPG strips (GE Healthcare) by swelling overnight. After focusing, the peptides were passively eluted into 72 contiguous fractions with MilliQ water/ 35% acetonitrile (CAN)/ 35% ACN + 0.1% formic acid (FA) using an in-house constructed IPG extraction robot (GE Healthcare Bio-Sciences AB, prototype instrument) into a 96-well plate (V-bottom, Greiner product #651201), which were then dried in a SpeedVac. The resulting fractions were dried and kept at −20°C.

### LC-MS/MS runs of the HiRIEF fractions

Online LC-MS was performed using a Dionex UltiMate™ 3000 RSLCnano System coupled to a Q-Exactive HF mass spectrometer (Thermo Fisher Scientific). Each fraction was subjected to MS analysis. Samples were trapped on a C18 guard-desalting column (Acclaim PepMap 100, 75μm x 2 cm, nanoViper, C18, 5 μm, 100Å), and separated on a 50 cm long C18 column (Easy spray PepMap RSLC, C18, 2 μm, 100Å, 75 μm x 50 cm). The nano capillary solvent A was 95% water, 5% DMSO, 0.1% formic acid; and solvent B was 5% water, 5% DMSO, 95% acetonitrile, 0.1% formic acid. At a constant flow of 0.25 μl min−1, the curved gradient went from 2% B up to 40% B in each fraction as shown in the Supplementary Data 6, followed by a steep increase to 100% B in 5 min. FTMS master scans with 60,000 resolution (and mass range 300-1500 m/z) were followed by data-dependent MS/MS (35 000 resolution) on the top 5 ions using higher energy collision dissociation (HCD) at 30% normalized collision energy. Precursors were isolated with a 1.2 m/z window. Automatic gain control (AGC) targets were 1E6 for MS1 and 1E5 for MS2. Maximum injection times were 100 ms for MS1 and 100 ms for MS2. Dynamic exclusion was set to 30 s duration. Precursors with unassigned charge state or charge state 1 were excluded. An underfill ratio of 1% was used.

### Analysis of LC-MS/MS runs

Orbitrap raw MS/MS files were converted to mzML format using msConvert from the ProteoWizard tool suite ^55^. Spectra were then searched using MSGF+ (v10072) ^56^ and Percolator (v2.08) ^57^, where search results from 8 subsequent fractions were grouped for Percolator target/decoy analysis. All searches were done against the human protein subset of Ensembl 99 in the Galaxy platform ^58^. MSGF+ settings included precursor mass tolerance of 10 ppm, fully-tryptic peptides, maximum peptide length of 50 amino acids and a maximum charge of 6. Fixed modifications were TMTpro 16-plex on lysines and peptide N-termini, and carbamidomethylation on cysteine residues, and a variable modification was used for oxidation on methionine residues. Quantification of TMTpro 16-plex reporter ions was done using IsobaricAnalyzer (v2.0) of the OpenMS project ^59^. PSMs found at 1% FDR (false discovery rate) were used to infer gene identities. Protein quantification by TMTpro 16-plex reporter ions was calculated using TMT PSM ratios. The median PSM TMT reporter ratio from peptides unique to a gene symbol was used for quantification. Protein false discovery rates were calculated using the picked-FDR method using gene symbols as protein groups and limited to 1% FDR ^60^.

### Data pre-processing and proteoform detection

Quantitative reporter ion signal for peptide spectrum matches was summarized on peptide level by summation. Reporter ion signals of all individual temperatures were normalized using the variance stabilizing normalization (vsn) ^61^ and converted to fold changes relative to the first temperature. Next, in order to assign similarly melting peptides found to map to a certain gene symbol, a graph for each gene symbol was created connecting all peptides (vertices) with weights (edges) corresponding to their similarity in melting profile. The similarity was computed with:

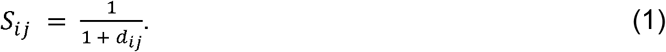

Where *d*_*ij*_ is the weighted Euclidean distance between two peptides across all cell lines:

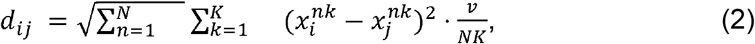

where 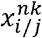 represents the fold change of peptide and respectively in cell line n at temperature k and v represents the number of valid comparison, i.e. 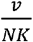 is the fraction of fold changes without missing values of either peptide. Obtained graphs were then used for community detection using the Leiden algorithm ^28^, however only gene symbols for which at least 10 peptides were identified and with at least two peptides per sample were subjected to this analysis (a detected community had to be supported by at least 5 peptides to be accepted). Peptides mapping to gene symbols for which these criteria were not fulfilled were grouped to single proteoforms, peptides mapping to gene symbols which were included in the community detection were assigned to proteoform groups if the modularity of the detected communities was higher than 0. Modularity was computed using the function *modularity()* of the igraph R package ^62^. Through the assignment of peptides to communities, proteoforms for each gene symbol were created. Summarization on proteoform level was performed by summation of raw peptide data assigned to individual communities, i.e., prior to vsn normalization. Obtained proteoform signal intensities were then normalized per temperature using vsn and relative fold changes to the lowest measured temperature were formed.

### Differential melting curve analysis

All proteoforms detected in at least 10 cell lines were fitted by a sigmoid function for each cell line individually. The sigmoid was fit using the NPARC R package implementation which is defined as:

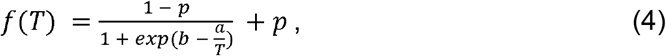

where *T* represents the temperature, *p* the plateau, and *a* and *b* are parameters affecting the slope and inflection point of the curve ^32,63^. Fits for individual cell lines (alternative model for the NPARC method) were accepted if they had a residual standard deviation of *s*_*res*_ < 0.1. The residual sum of squares (RSS) was computed across cell lines as: 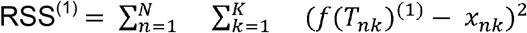 and melting points (*f*(*T*_*m*_) = 0.5) and areas under the melting curve were computed for accepted fits of cell line-specific proteoform thermal profiles. Null models were fit using the same sigmoid model (4) for each proteoform across all cell lines for which an alternative model fit was accepted. The null model RSS was computed as: 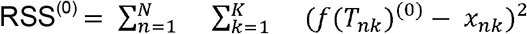. Based on the RSS of both models, an *F*-statistic was computed with:

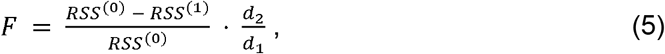

where the degrees of freedom *d*_1_ = *v*_1–_*v*_0_ and *d*_*2*_ = *p*_*i* –_*v*_0_, with *p*_*i*_, *v*0 and v1representing the number of observations for protein i, the number of parameters of the null and alternative model, respectively ^32^. Proteoforms with an *F*-statistic above the 90% percentile were considered significantly different across cell lines.

### Differential proteoform-proteoform coaggregation analysis

In order to test for pairs of proteoforms which coaggregated in some cell lines, but melted differentially in others, we adapted our previous approach for testing this between two conditions ^40^. We started by extending the list of highly confident string interactions (combined score >= 950) by all possible proteoform interactions, i.e. if protein A was previously annotated to interact with protein B and we detected three proteoforms for protein A and two for B, we replaced this entry by all possible 3 × 2 combinations. Next, we tested for coaggregation of pairs of proteoforms in all individual cell lines using the approach described by ^19^. All pairs of proteoforms which showed significant coaggregation (*p*_adj._ < 0.1) in at least one of the cell lines were included for the differential analysis across cell lines. The test statistic for differences in coaggregation across cell lines was determined by computing 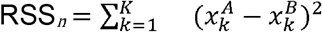 across all temperatures *K*, between all annotated pairs of proteoforms *A* and *B* for all individual cell lines *n*, ranking all RSS_*n*_ and computing:

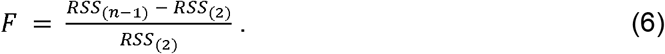

Above, RSS_(*n*-*1*)_ and RSS_(*2*)_ represent the second highest and the second lowest RSS. In this way the *F*-statistic became only large for cases in which at least two cell lines featured small and big differences between the melting curves of the two proteoforms, respectively. We considered *F*-statistics higher than the 90% percentile as significant.

### Proteoform thermal stability and drug sensitivity correlation analysis

Proteoform thermal stabilities were associated with drug sensitivity scores (DSS) by performing correlation analyses between the area under the melting curves of proteoforms found to differ across cell lines (90% percentile of observed *F*-statistics obtained from the NPARC analysis) with the DSS of the respective cell lines for all drugs with a minimal effect (DSS < 6 for at least one cell line) using the R package *limma* ^*46,64*^. Results obtained for all proteoforms and drugs were jointly adjusted for multiple testing using the method by Benjamini and Hochberg ^65^. Proteoform-DSS associations with an adjusted p-value of less than 0.1 were considered significant.

### Differential RNA-seq analysis

Differential RNA-seq analysis was performed using the *DESeq2* ^66^. The sex of the cell line donors was included as a covariate in the design formula, when testing for a difference in conditions.

### Gene set enrichment analysis

Gene set enrichment analysis was performed using the log fold changes computed between the conditions of all genes using the R Bioconductor package *fgsea* ^67^.

### Gene Ontology enrichment

Gene Ontology enrichment was performed using the R Bioconductor package *clusterprofiler* ^*68*^.

### CETSA temperature range analysis

COG-355 and ALL-PO cells were cultured as previously described. Cell suspensions were then centrifuged at 300g for 5 min, the supernatant media was discarded and the cells were washed twice with Hank’s Balanced Salt Solution (HBSS, Gibco/Life Technologies). Pelleted cells were resuspended in HBSS and 75μL cell suspension were aliquoted to 0.2 mL tubes. Samples were then heated in a temperature range of 37-70°C in a Veriti Thermal Cycler (Applied Biosystems/Thermo Fisher Scientific) for 3 min, followed by 3 min cooling at room temperature and immediate snap-freezing in liquid nitrogen. The cells were then lysed by three repeated freeze-thawing cycles and centrifuged at 21000 x *g* for 40 min at 4□. The cleared supernatants were transferred to new tubes, denatured in LDS sample buffer (Thermo Fisher Scientific) and analyzed by western blotting.

### Western blotting

Cleared protein supernatants were denatured in LDS sample buffer (Thermo Fisher Scientific), resolved by SDS-PAGE using NuPAGE 4 to 12%, Bis-Tris Gel (Invitrogen, Thermo Fisher Scientific) and NuPAGE MES SDS Running Buffer (Invitrogen, Thermo Fisher Scientific), and transferred to Nitrocellulose membranes (Invitrogen, Thermo Fisher Scientific). SeeBlue Plus2 Pre-stained Protein Standard was used as protein ladder (Invitrogen, Thermo Fisher Scientific). Afterward, the membranes were blocked with 5% non-fat dry milk in TBST (Thermo Fisher Scientific) and incubated with primary antibodies for the appropriate target. TMPO/LAP2 (Thermo Fisher Scientific, cat. No; A304-838A-M 1:1000 dilution), PSAP (Thermo Fisher Scientific, cat# PA5-21340, 1:1000 dilution) and Saposin-C (Santa Cruz Biotechnology cat# sc-374119, 1:500 dilution) antibodies were used for western blot to detect corresponding targets. Following overnight primary incubation at 4°C, blots were rinsed using TBST and incubated with the appropriate horseradish peroxidase (HRP)-conjugated secondary antibodies (Millipore, cat no. AP127P for mouse primary ab and Santa Cruz Biotechnology cat# sc-2005) for rabbit primary ab, both used at a dilution of 1:5000). All antibody incubations were diluted in 5% non-fat dry milk in TBST. Protein bands were developed with Clarity ECL Substrate Chemiluminescent HRP substrate (Bio-Rad) in a iBright CL1000 Imaging System (Invitrogen, Thermo Fisher Scientific). Bands were quantified using iBright Analysis Software version 4.0.1 (Thermo Fisher Scientific). Images of the full uncropped blots are provided with annotation (Supplementary Figure 12).

## Data availability

All proteomics datasets have been deposited on PRIDE. Images of the full uncropped blots corresponding to Figure 2d and Supplementary Figure 5c are provided with annotation in Supplementary Figure 12.

## Code availability

All code used to perform the computational analyses described and to reproduce the figures is available at: https://github.com/nkurzaw/deepPedAllMeltome

## Acknowledgements

This study was supported by grants from the Swedish Childhood Cancer Foundation (R.J., grant reference TJ2016-0035, PR2016-0019 and PR2019-0025; M.S., TJ2019-0023), the Swedish Research Council (R.J., grant reference 2017-01653), Felix Mindus Contribution to Leukemia research (R.J.), Dr Åke Olsson Foundation for Hematological Research (R.J., grant reference 2017-00437 and 2021-00130), Cancer Society Stockholm and the King Gustaf V Jubilee Fund (R.J., grant references 174182, 194111 and 204092), Magnus Bergvalls Stiftelse (R.J., grant reference 2017-02421 and 2016-01841). R.J. acknowledge Karolinska Institutet and SciLifeLab. N.K. was supported by a fellowship of the EMBL International PhD programme. A.M. was supported by a fellowship from the EMBL Interdisciplinary Postdoc (EI3POD) programme under Marie Skłodowska-Curie Actions COFUND (grant number 664726).

## Author contributions

R.J. conceived and coordinated the study and acquired funding. M.M.S. co-supervised the analysis with A.M. and provided resources and funding together with W.H.. E.K. and R.J. performed the LC-MS experiments with support from A.A. and G.M.. N.K. and M.S. developed and performed the proteoform detection and analysis. I.L. and I.B. performed validation experiments. N.K. wrote the manuscript with M.S., A.M., M.M.S. and R.J.. All authors contributed to finalizing the manuscript and approved the final version.

