## Supplementary Figures for "Deep thermal proteome profiling for detection of proteoforms and drug sensitivity biomarkers"

Supplementary Figures 1-12 related to:

### Deep thermal proteome profiling for detection of proteoforms and drug sensitivity biomarkers

Kurzawa, Stahl et al.

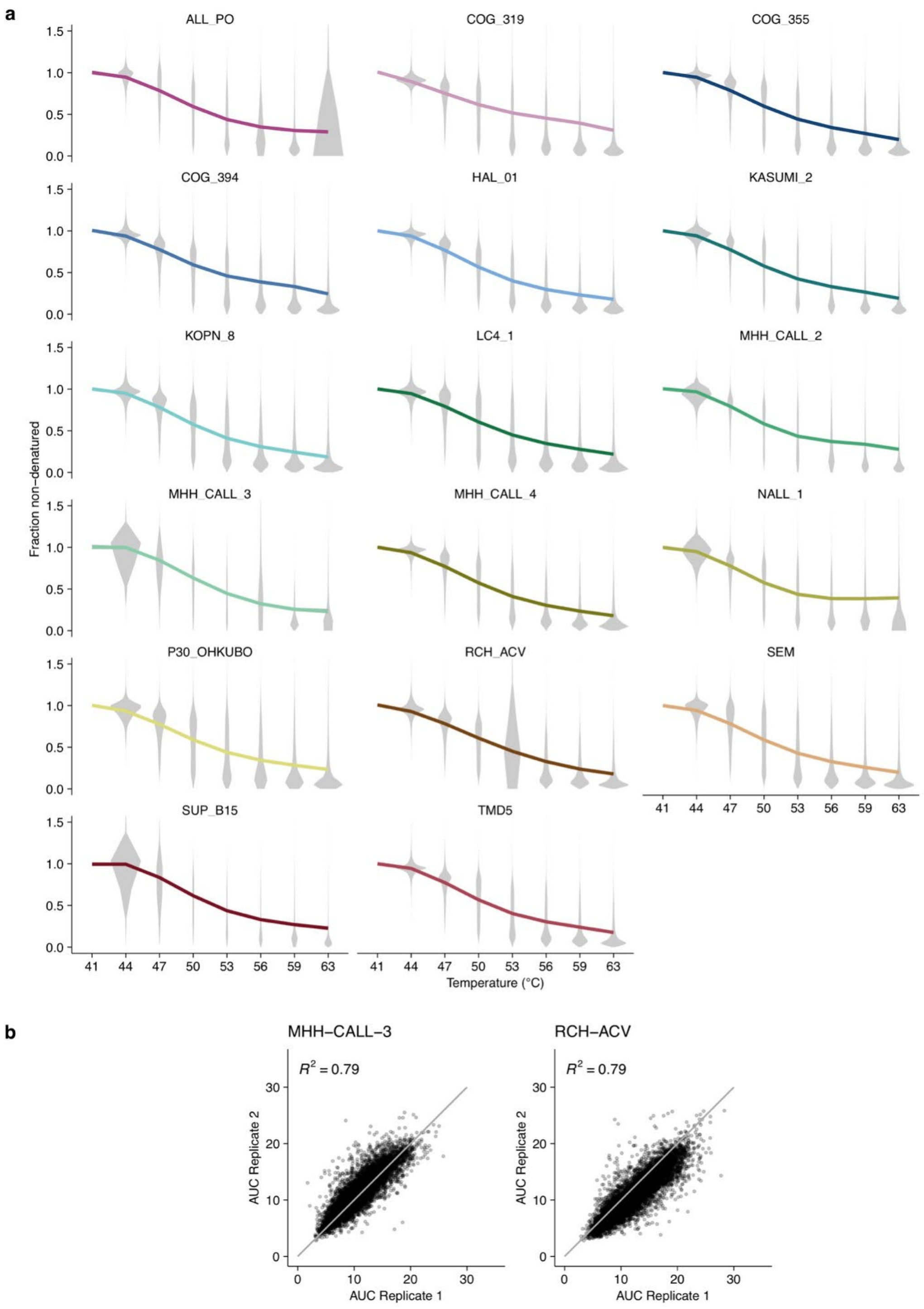


**Supplementary Figure 1**: Global melting profiles of cell lines. a) Average melting profiles after normalization across all peptides identified and quantified in all cell lines not shown in the main. b) Scatterplots of area under the melting curves (AUC) obtained from two biological replicates of MHH-CALL-3 and RCH-ACV.

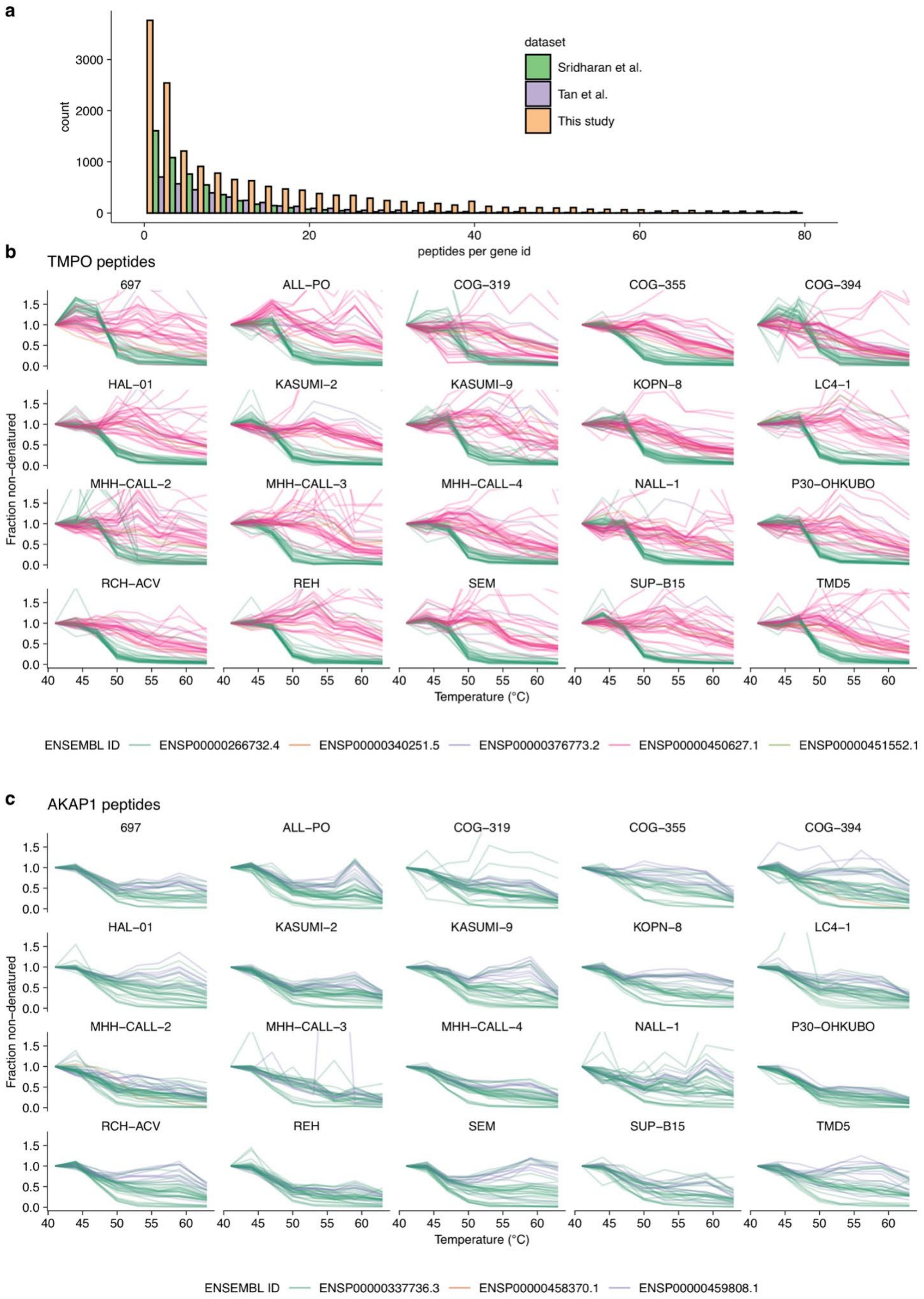


**Supplementary Figure 2**: Gene symbol level peptide analysis. a) Comparison of peptides mapping to gene symbols in this and previous studies. b, c) Differential melting profiles of peptides mapping to gene symbols resembling annotated proteoforms.

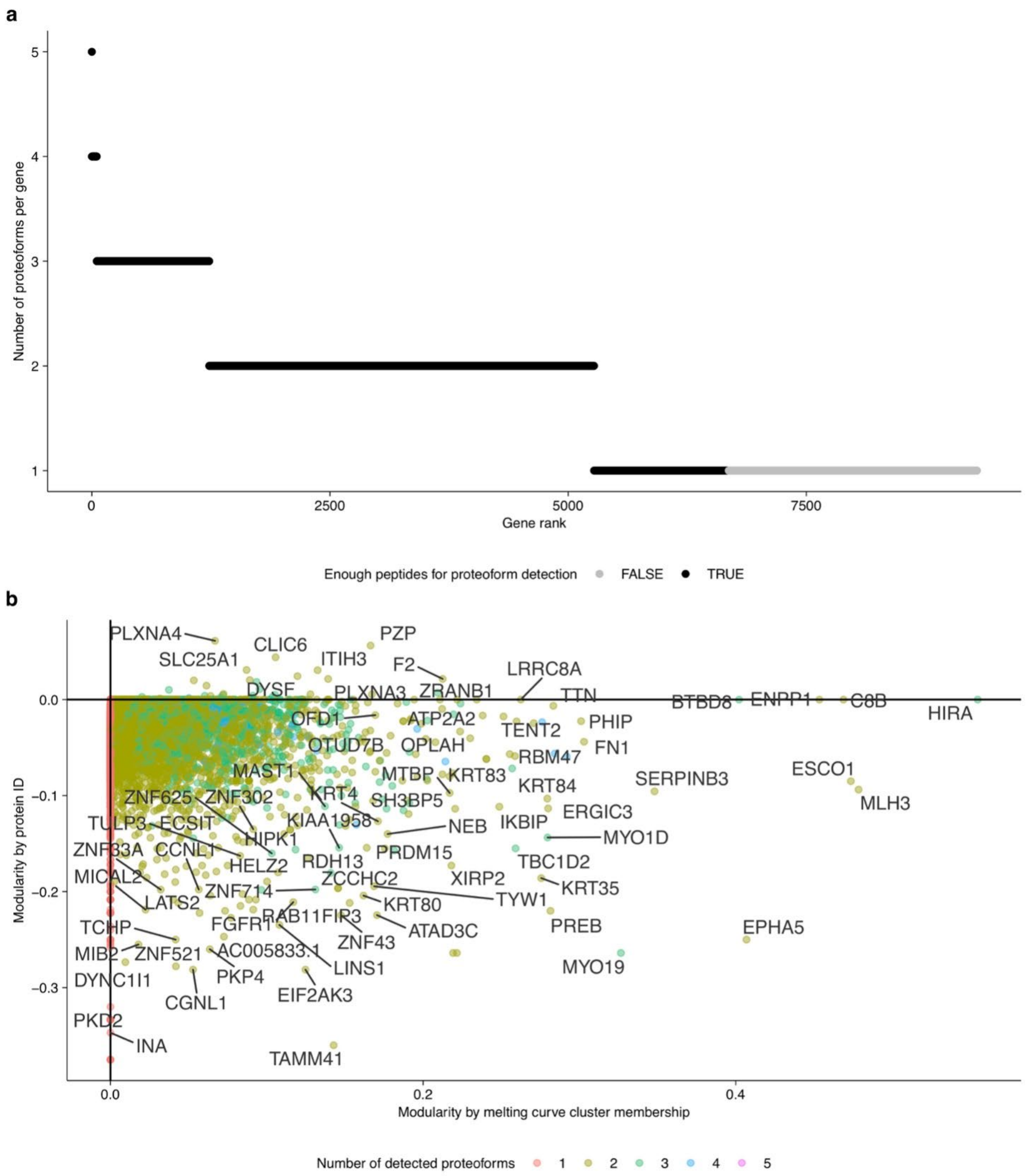


**Supplementary Figure 3**: Proteoform detection statistics. a) Dotplot of the number of detected proteoforms per gene symbol. b) Scatterplot of modularity by protein ID (ENSEMBL) versus modularity by detected proteoform based on melting curve similarity.

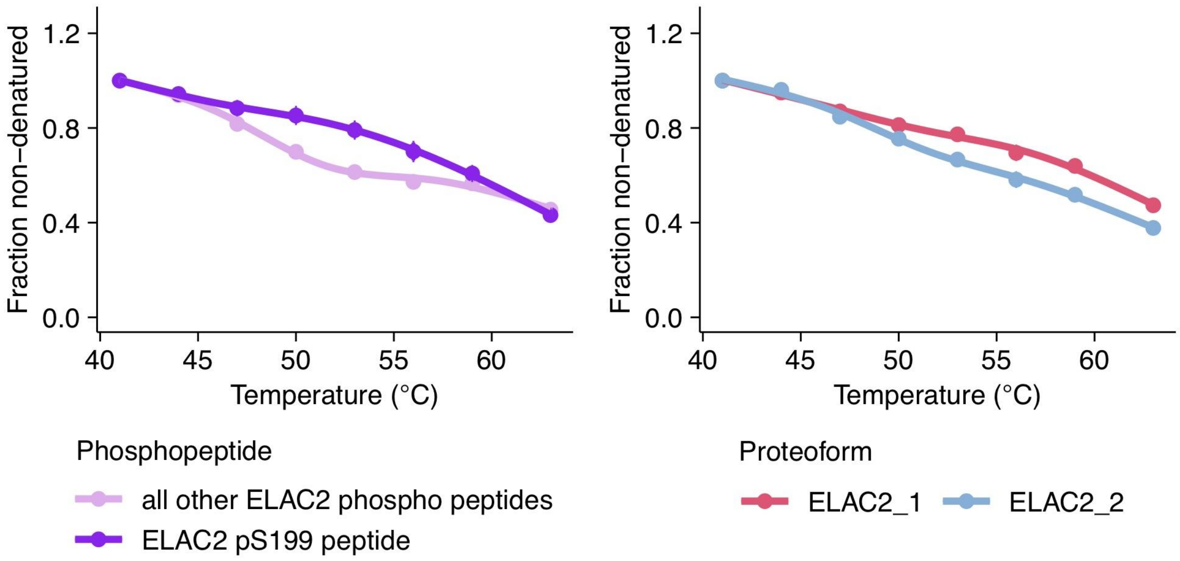


**Supplementary Figure 4**: ELAC2 pS199 phosphopeptide melting. a) Average melting profiles across cell lines for the ELAC2 peptide phosphorylated on serine 199 versus all other phosphopeptides found for ELAC2. b) Average melting profiles across cell lines for the two ELAC2 proteoforms.

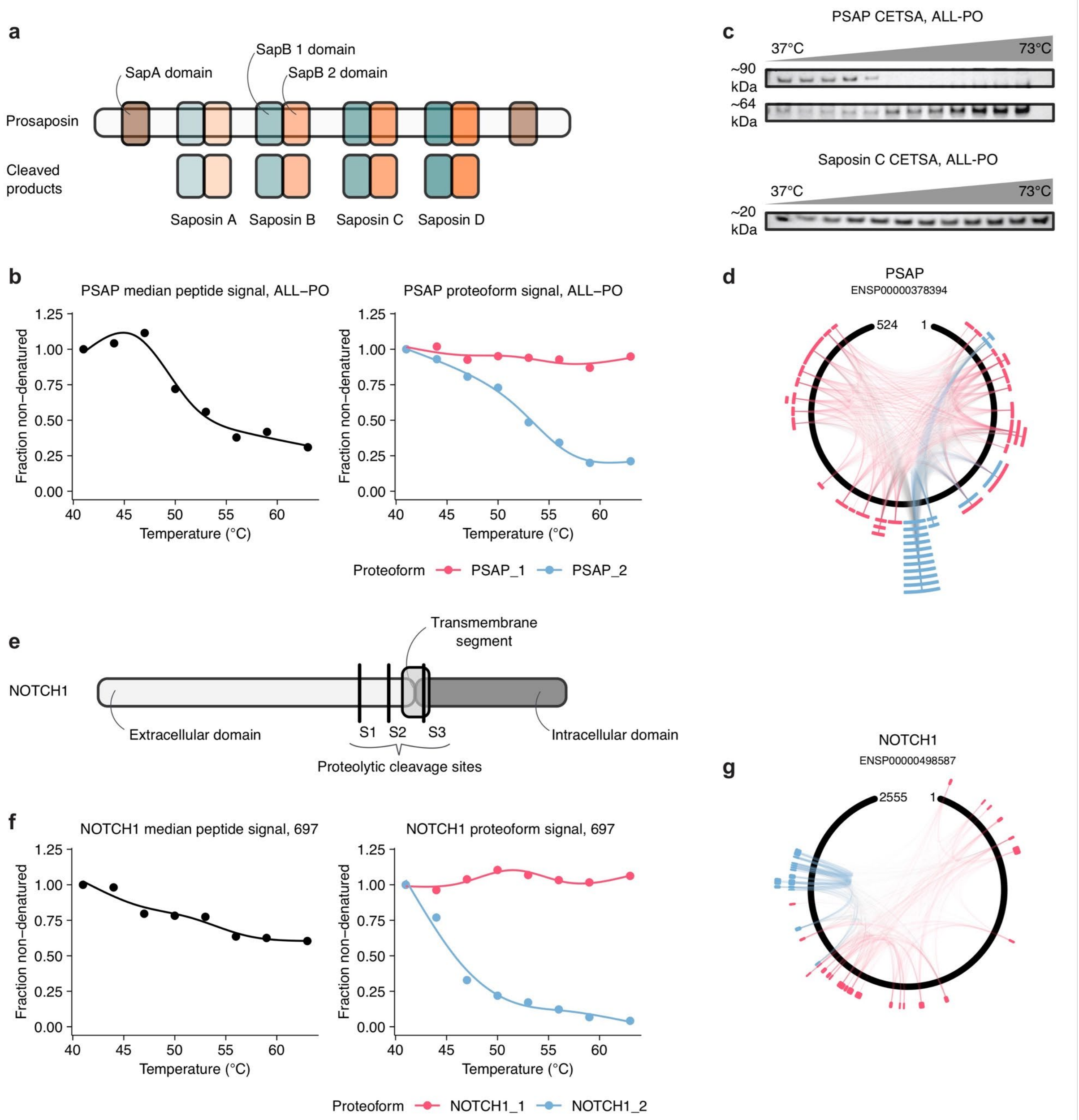


**Supplementary Figure 5**: Proteoform examples indicative of proteolytic cleavage detected by similarity of peptide melting profiles. a) Schematic of the protein domains of Pre-saposin (PSAP) and its cleaved products. PSAP is a protein which exists as an integral membrane protein, but also is proteolytically processed to form Saposin A, B, C and D (Kishimoto et al., 1992). b) Melting curves of the median peptide signal per PSAP gene symbol (left) and by detected proteoforms. c) A multi-band signal with differential thermal stabilities was also observed when performing a CETSA experiment with a PSAP antibody, whereas when using a Saposin C-specific antibody, solely a high thermal stability signal could be detected. When considering the mapping of the proteoform-specific peptides to the protein sequence (d), it became apparent that PSAP_1 likely reflected the non-cleaved form of PSAP. The peptides of PSAP_2, which featured lower thermal stability than those of PSAP_1,  mapped to the second SapB domain and N-terminal of the protein, suggesting that this proteoform could capture an intermediate cleavage product, since it is known that the processed Saposins exhibit high thermal stability (Kishimoto et al., 1992) which is also reflected by the CETSA signal of Saposin C. e) Schematic of the protein domains of the transmembrane receptor NOTCH1. Upon ligand binding NOTCH1 releases a cleaved C-terminal domain which then functions as a transcriptional effector. f) Melting curves of the median peptide signal per NOTCH1 gene symbol (left) and by detected proteoforms. Considering the mapping of peptides assigned to the different proteoforms (g) revealed that NOTCH1_1 likely represented the membrane bound pool of the protein constituting high thermal stability, whereas NOTCH1_2 likely reflected the intracellular cleavage product.

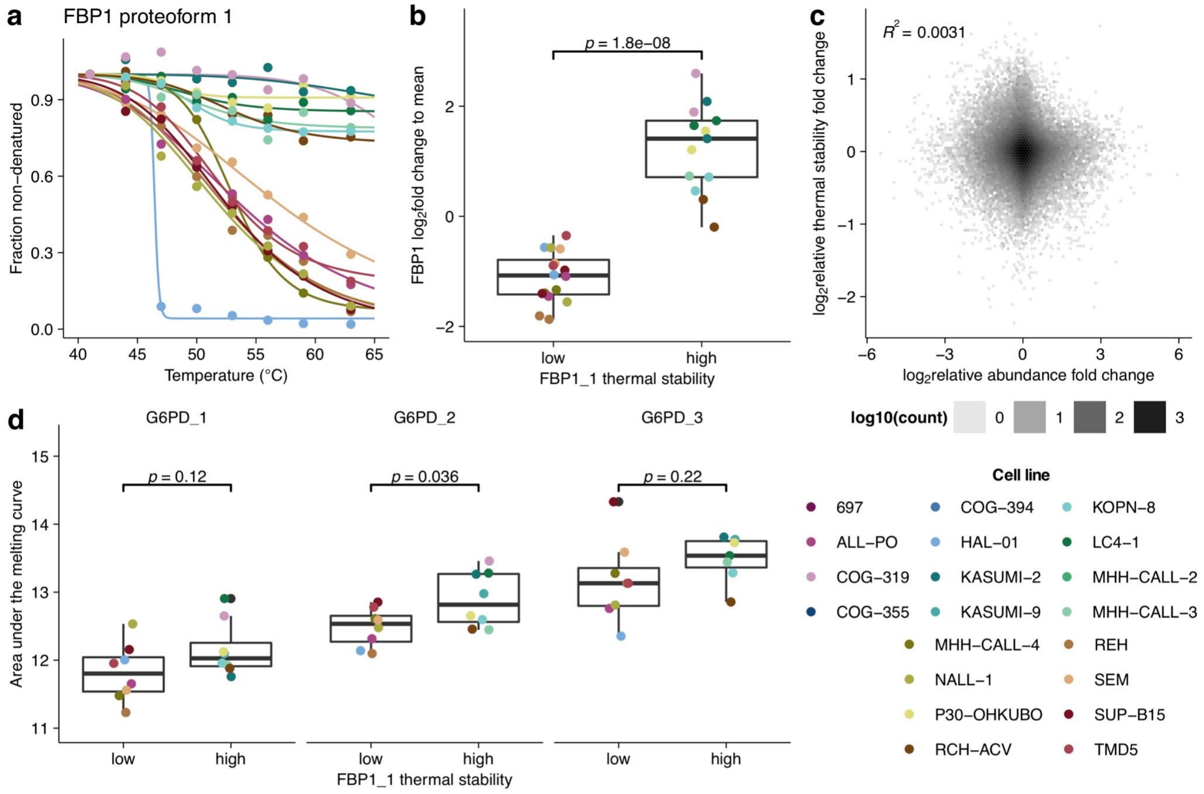


**Supplementary Figure 6**: Thermal stability and abundance of FBP1. a) Melting profiles of FBP1 proteoform 1 (FBP1_1) in the different cell lines. b) Boxplot of FBP1 $\log_2$ abundance fold change over mean in cell lines with low and high FBP1_1 thermal stability. c) Global correlation of $\log_2$ thermal stability fold change over mean and $\log_2$ abundance fold change over mean across the cell lines. d) Boxplots of the $\log_2$ abundance fold changes over the mean of all G6PD proteoforms in cell lines with low and high FBP1_1 thermal stability. Center lines in all box plots represent the median, the bounds of the boxes are the 75 and 25% percentiles i.e., the interquartile range (IQR) and the whiskers correspond to the highest or lowest respective value or if the highest value is an outlier (greater than 1.5 * IQR from the bounds of the boxes) it is exactly 1.5 * IQR.

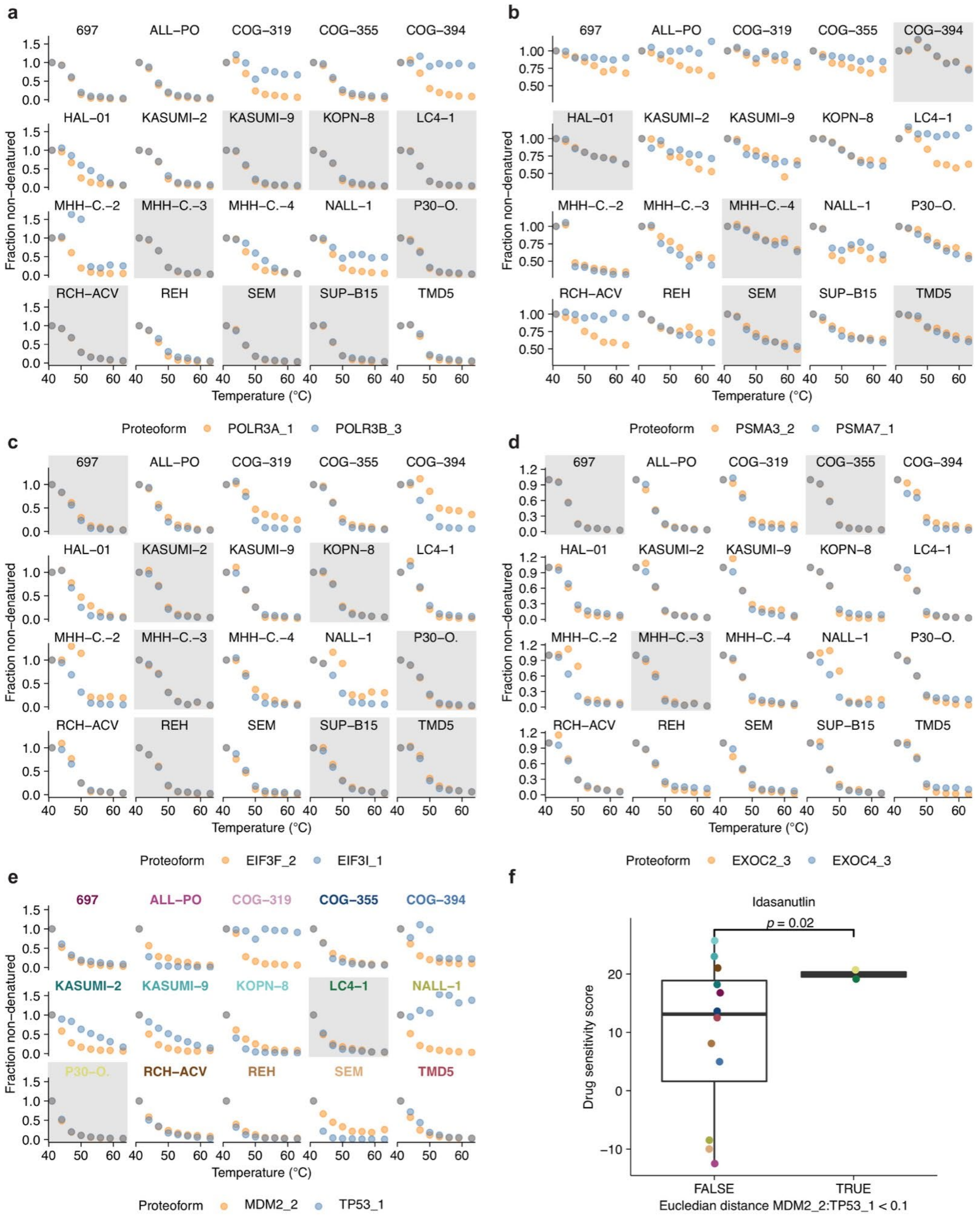


**Supplementary Figure 7**: Additional profiles of pairs of proteoforms found to coaggregate differentially across cell lines. Melting profiles of the differentially coaggregating proteoform pairs a) POLR3A_1 and POLR3B_3, b) PSMA3_2 and PSMA7_1, c) EIF3F_2 and EIF3I_1, d) EXOC2_3 and EXOC4_3 and e) MDM2_2 and TP53_1. Profiles indicative of coaggregation (Euclidean distance between profiles < 0.1) are shown with a gray background and otherwise with a white background. f) Boxplot of drug sensitivity scores for Idasanutlin for cell lines which do or do not show coaggregation of MDM2_2 and TP53_1. Center lines in all box plots represent the median, the bounds of the boxes are the 75 and 25% percentiles i.e., the interquartile range (IQR) and the whiskers correspond to the highest or lowest respective value.

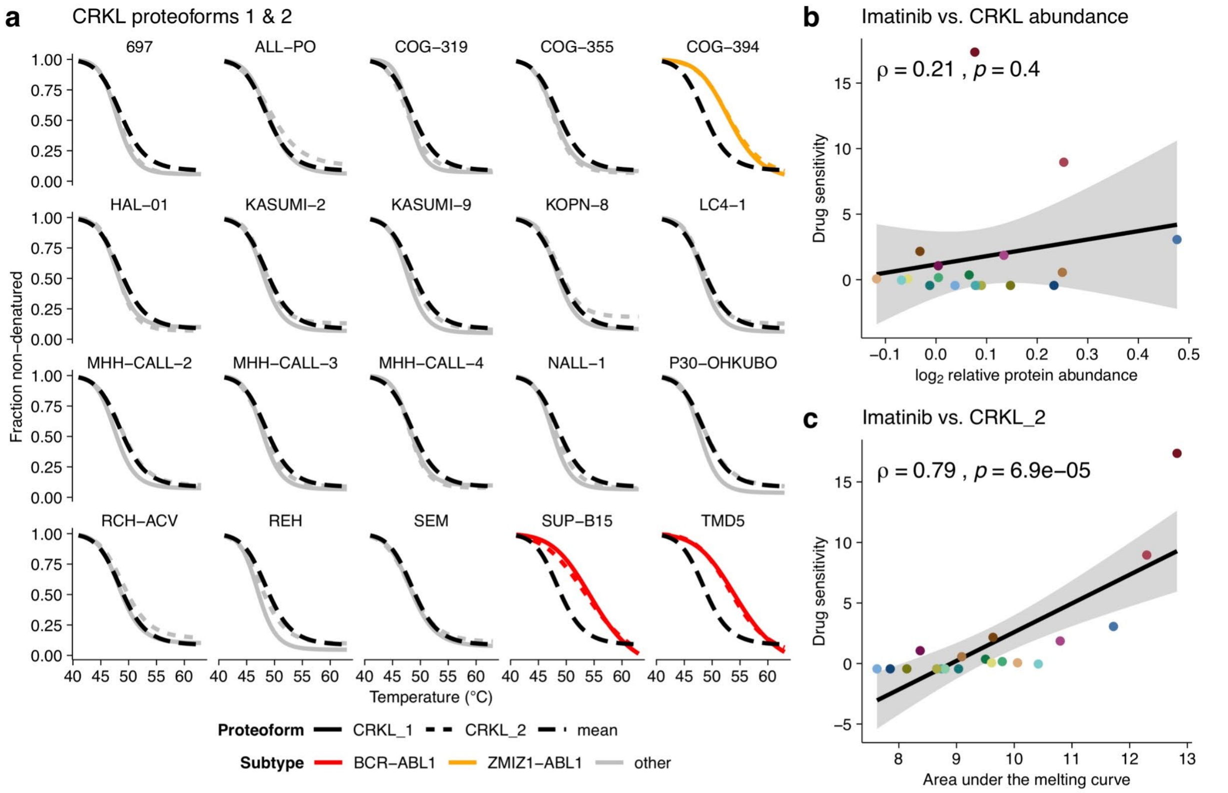


**Supplementary Figure 8**: Correlation of CRKL thermal stability and abundance with drug sensitivity. a) Melting profiles of CRKL proteoform 1 and 2 (CRKL_1 and CRKL_2) across cell lines. b) Correlation of CRKL abundance and sensitivity to imatinib. c) Correlation of CRKL_2 thermal stability and sensitivity to imatinib.

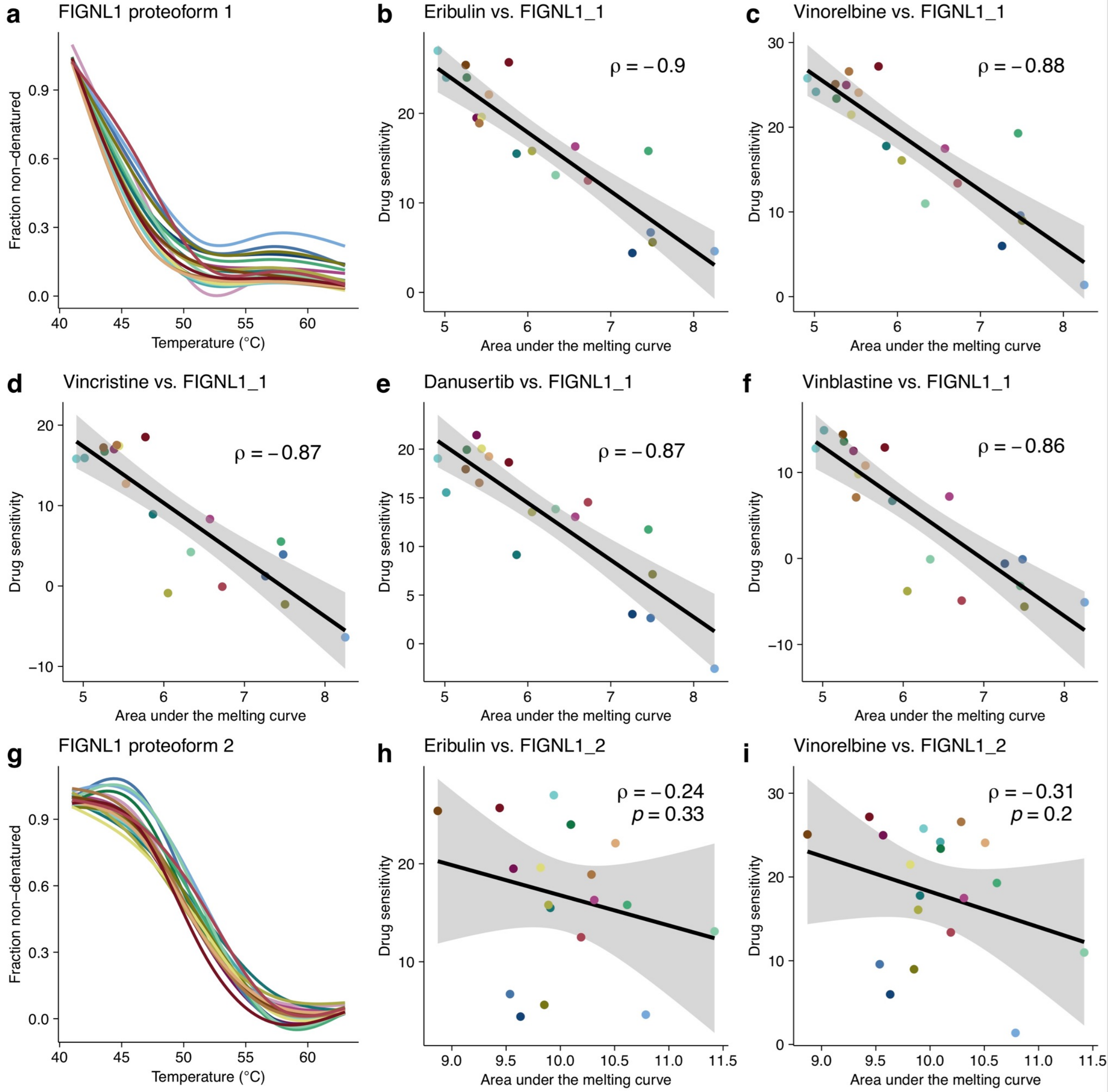


**Supplementary Figure 9**: Correlation of FIGNL1 thermal stability and abundance with drug sensitivity. a) Melting profiles of FIGNL1 proteoform 1 (FIGNL1_1) across cell lines. Significant FIGNL1_1 thermal stability association with drug sensitivity to b) eribulin, c) vinorelbine, d) vincristine, e) danusertinib and f) vinblastine. g) Melting profiles of FIGNL1 proteoform 2 (FIGNL1_2) across cell lines. Correlation of FIGNL1_2 thermal stability and sensitivity to h) erubilin and i) vinorelbine.

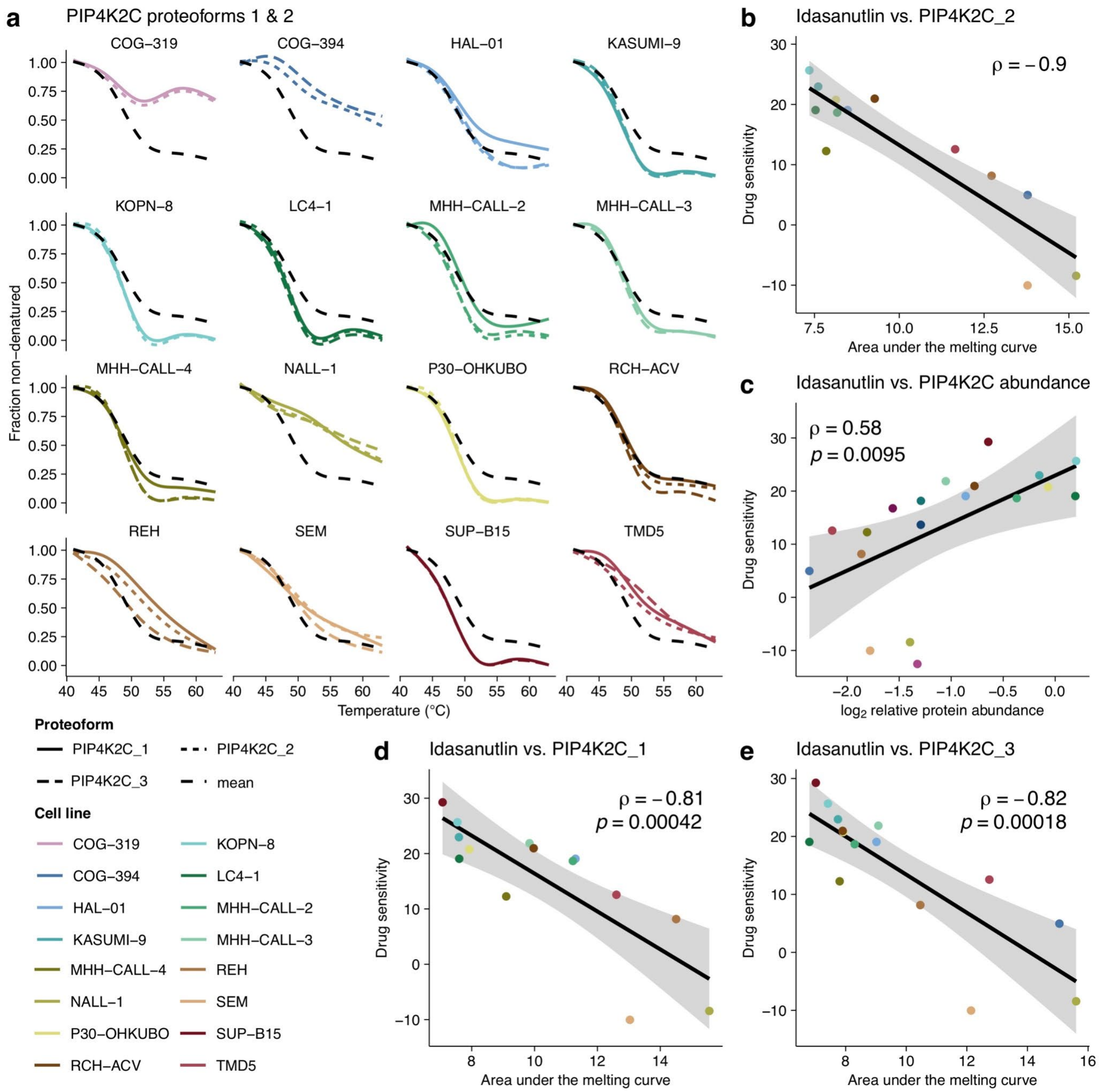


**Supplementary Figure 10**: Correlation of PIP4K2C thermal stability and abundance with drug sensitivity. a) Thermal stability profile of the PIP4K2C proteoforms. b) Scatterplot of sensitivity to idasanulin versus PIP4K2C_2 thermal stability. c) Correlation of idasanulin sensitivity versus PIP4K2C $\log_2$ abundance fold changes over the mean across cell lines. Scatterplots of sensitivity to idasanulin versus d) PIP4K2C_1 and e) PIP4K2C_3 thermal stability.

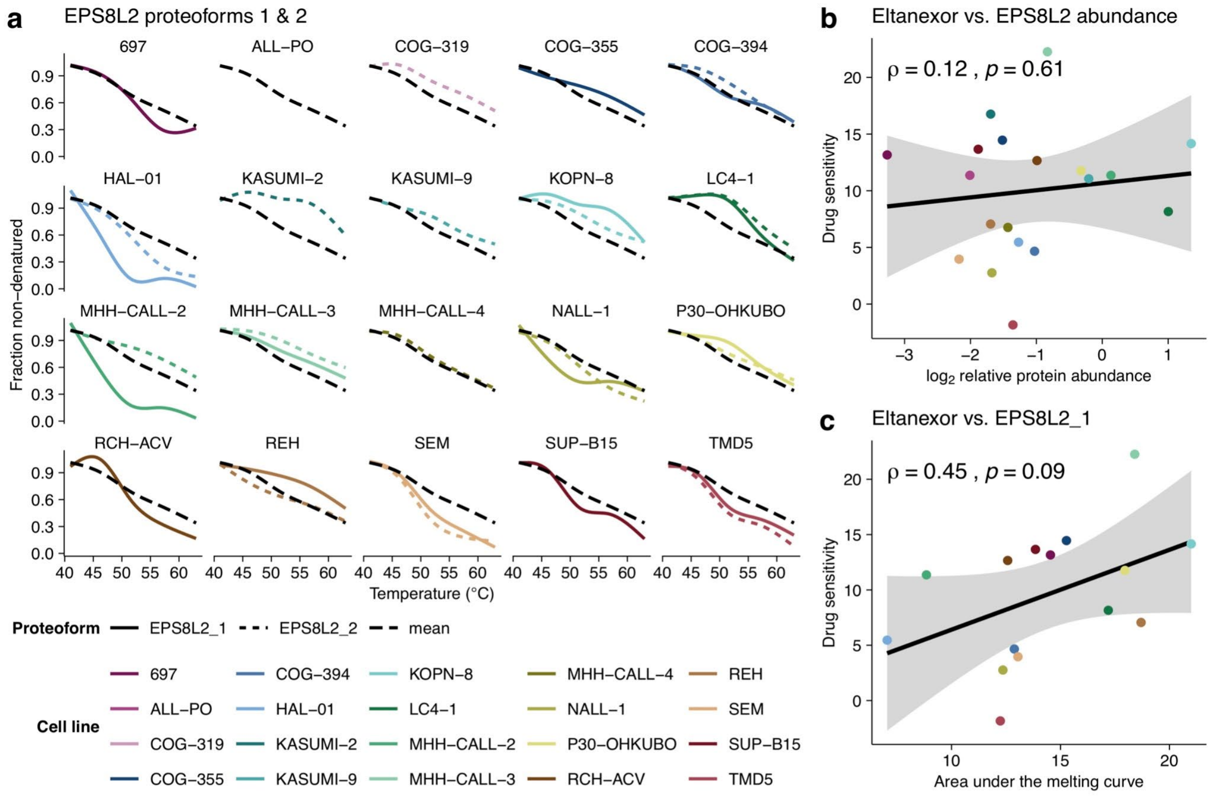


**Supplementary Figure 11**: Correlation of EPS8L2 thermal stability and abundance with drug sensitivity. a) Thermal stability profile of the EPS8L2 proteoforms. b) Scatterplot of sensitivity to Eltanexor versus EPS8L2 $\log_2$ abundance fold changes over the mean across cell lines. c) Scatterplot of sensitivity to Eltanexor versus EPS8L2_1 thermal stability across cell lines.

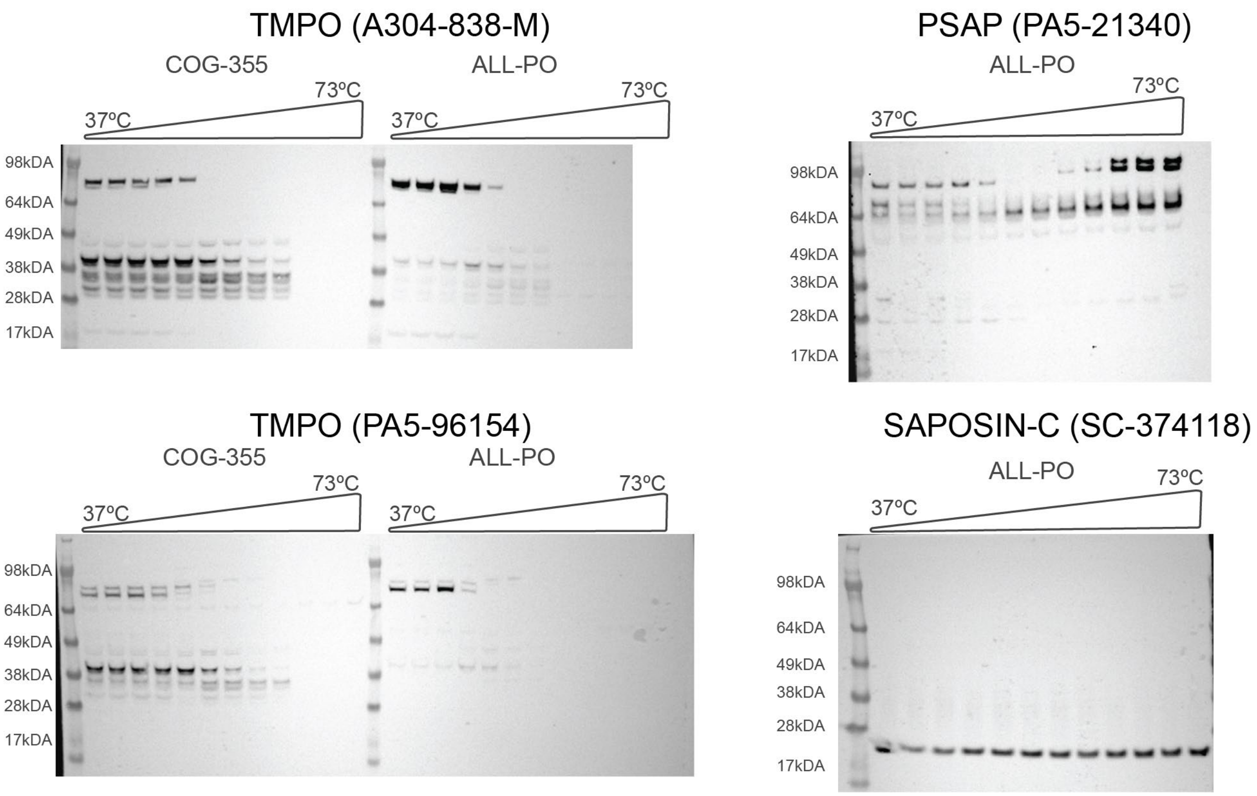


**Supplementary Figure 12**: Annotated raw western blot images from the CETSA experiments for TMPO in COG-355 and ALL-PO cell lines and PSAP in ALL-PO cell line. The CETSA temperature range (37-73℃) was performed once (n = 1) for each cell line. Bands for molecular weight markers (M.W., SeeBlue™ Plus2) are annotated in each panel. The catalog nr for each antibody used is shown in brackets for each protein.
